# Population genomics and antimicrobial resistance in *Corynebacterium diphtheriae*

**DOI:** 10.1101/2020.05.19.101030

**Authors:** Melanie Hennart, Leonardo G. Panunzi, Carla Rodrigues, Quentin Gaday, Sarah L. Baines, Marina Barros-Pinkelnig, Annick Carmi-Leroy, Melody Dazas, Anne-Marie Wehenkel, Xavier Didelot, Julie Toubiana, Edgar Badell, Sylvain Brisse

## Abstract

*Corynebacterium diphtheriae*, the agent of diphtheria, is a genetically diverse bacterial species. Although antimicrobial resistance has emerged against several drugs including first-line penicillin, the genomic determinants and population dynamics of resistance are largely unknown for this neglected human pathogen.

Here we analyzed the associations of antimicrobial susceptibility phenotypes, diphtheria toxin production and genomic features in *C. diphtheriae.* We used 247 strains collected over several decades in multiple world regions, including the 163 clinical isolates collected prospectively from 2008 to 2017 in France mainland and overseas territories.

Phylogenetic analysis revealed multiple deep-branching sublineages, grouped into a Mitis lineage strongly associated with diphtheria toxin production, and a *tox*-negative Gravis lineage with few *tox*^+^ exceptions including the 1990s ex-Soviet Union outbreak strain. The distribution of susceptibility phenotypes allowed proposing ecological cutoffs for most of the 19 agents tested, thereby defining acquired antimicrobial resistance. Penicillin resistance was found in 17.2% of prospective isolates. Four isolates were multidrug resistant (>8 agents), including to penicillin and macrolides. Homologous recombination was frequent (r/m = 5) and horizontal gene transfer contributed to the emergence of antimicrobial resistance in multiple sublineages. Genome-wide association mapping uncovered genetic factors of resistance, including an accessory penicillin-binding protein (PBP2m) located in diverse genomic contexts. Gene *pbp2m* is widespread in other *Corynebacterium* species and its expression in *C. glutamicum* demonstrated its effect against several beta-lactams. A novel 73-kb *C. diphtheriae* multi-resistance plasmid was discovered.

This work uncovers the dynamics of antimicrobial resistance in *C. diphtheriae* in the context of phylogenetic structure, biovar and diphtheria toxin production, and provides a blueprint to analyze re-emerging diphtheria.

## INTRODUCTION

Diphtheria, if untreated, is one of the most severe bacterial infections of humans. It typically affects the upper respiratory tract causing pseudomembrane formation, sometimes leading to suffocation and death. The infection can be complicated by toxinic symptoms, caused by the diphtheria toxin. Other forms of disease are skin and invasive infections, including endocarditis ^1,2^.

The agent of diphtheria is *Corynebacterium diphtheriae*, a member of the phylum Actinomycetes ^34^. The diphtheria toxin, encoded by the *tox* gene, is carried by lysogenized corynephages within the chromosome of some *C. diphtheriae* strains ^5,6^. Concern exists about the possibility of lysogenic conversion of previously non-toxigenic strains during colonization, infection or transmission chains ^7^. However, knowledge on the microevolutionary dynamics between *tox*-positive and *tox*-negative strains is limited. The high genetic diversity of *C. diphtheriae* strains underlies their variable colonization, adhesion and pathogenicity properties ^8–10^. Although three main biovars (Mitis, Gravis and Belfanti) are distinguished since the 1950s, their phylogenetic relationships are poorly defined ^11–13^.

Diphtheria used to be one of the deadliest infections in young children, but has been largely controlled by vaccination with the highly effective toxoid vaccine ^14^. Even so, thousands of cases of diphtheria are still reported annually ^15^, and large outbreaks can quickly follow the disruption of public health systems ^14,16–18^. In countries with high vaccination coverage, diphtheria cases are associated with travel and migration from endemic regions ^19–21^. As diphtheria vaccination is performed using an inactivated form of diphtheria toxin, it is not considered to prevent asymptomatic colonization and silent transmission of the pathogen, which still circulates and is the object of intense epidemiological surveillance ^4^. However, vaccine preparations may include other antigens and the impact of vaccination on *C. diphtheriae* evolution deserves further studies ^22^.

Clinical management of infections with toxigenic isolates includes treatment with diphtheria antitoxin (DAT), which can prevent or reduce toxinic complications ^4^. Nevertheless, antimicrobial treatment is critical in clinical management of both *tox*-positive and *tox*-negative infections, as it contributes to the elimination of the bacteria within the patient and limits transmission to novel individuals ^23^. With DAT production being threatened ^24^, antimicrobial treatment might become even more critical in diphtheria therapy.

Penicillin is the first-line therapeutics to treat diphtheria, with erythromycin being recommended in case of allergy ^25^. Both antimicrobial agents are effective for the treatment of diphtheria ^23,26^. However, reduced susceptibility or full resistance of *C. diphtheriae* to penicillin has been reported from multiple world regions ^27–31^. Resistance against other antimicrobial agents including erythromycin has also been reported ^23,26,27,32–35^. Although rare, multidrug resistant *C. diphtheriae* have been described ^26,27,32,35,36^.

Antimicrobial resistance genes have been described in *C. diphtheriae*, including the erythromycin resistance gene *ermX* on plasmid pNG2 ^37^ and genes *dfrA16, qacH* and *sul1* carried on a class 1 integron, mobilized by IS*6100* ^38^. However, the prevalence and phylogenetic distribution of resistance genes in *C. diphtheriae* clinical isolates are unknown. Six chromosomal penicillin-binding proteins (PBP) have been reported in *C. diphtheriae* ^39^, but so far no association between *pbp* or other genetic variation and penicillin resistance has been described. Understanding the genetic basis of antimicrobial resistance in *C. diphtheriae* would improve our ability to diagnose and track its spread.

The aims of this study were (i) to characterize antimicrobial resistance phenotypes in a large collection of *C. diphtheriae* strains with diverse geographical and temporal origins, and to uncover genomic determinants of resistance; and (ii) to analyze the population structure of *C. diphtheriae* and define associations between antimicrobial resistance, diphtheria toxin production, biovars and phylogenetic sublineages.

## RESULTS

### Provenance and microbiological characteristics of *C. diphtheriae* isolates

We studied 247 *C. diphtheriae* strains of diverse geographic and temporal origins (**Figure 1)**. This collection included 163 isolates prospectively collected between 2008 and 2017 from French mainland and overseas territories, 15 older (1981-2006) French clinical isolates, 65 ribotype reference strains ^40^ and 4 other reference strains. All isolates were confirmed as *C. diphtheriae* (excluding *C. belfantii* and *C. rouxii*) based on an average nucleotide identity (ANI) value higher than 96% with the *C. diphtheriae* type strain NCTC11397^⊤^.

**Figure 1.**
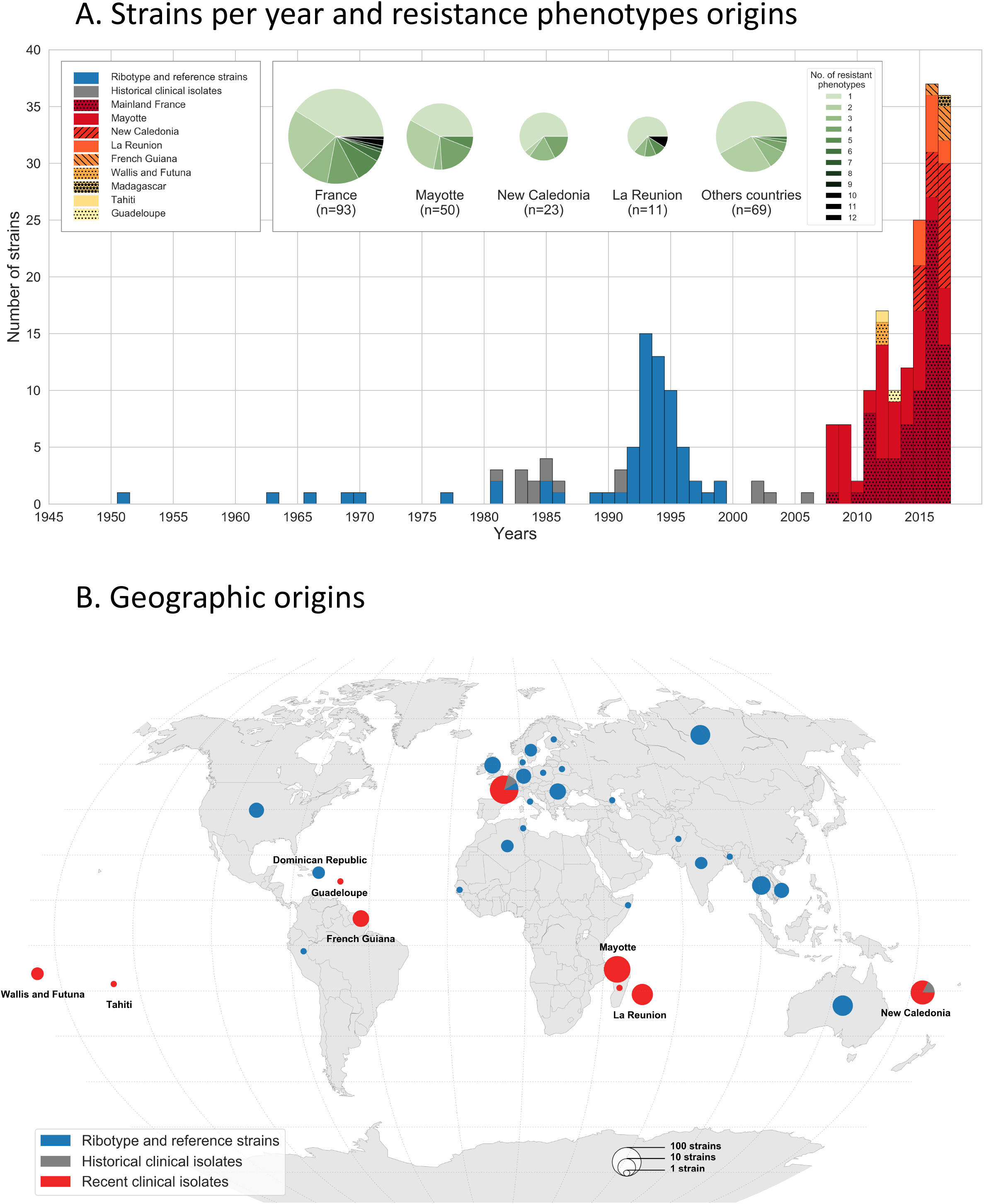
Temporal and geographical distribution of strains studied. A. Number of strains per year, 1945-2017. The 2008-2017 clinical isolates are represented in red or yellow shaded patterns (see key), whereas the older clinical isolates are in grey and the reference strains in blue. The inset shows pie charts with the frequency of resistance phenotypes among strains from the four most represented geographic origins; the remaining ones are pooled in the right-most pie chart. B. Geographic origins of strains from the three subsets.

Approximately one third (n = 78, 32%) of isolates were *tox*-positive (as defined by the detection of the *tox* gene by PCR), whereas the remaining 169 isolates (68%) were *tox*-negative. The proportions of *tox*-positive isolates were 42%, 34% and 2% among reference strains, 2008-2017 clinical isolates and older clinical isolates, respectively (**Figure S1**). Of the 78 *tox*-positive isolates, 17 (21.8%) were negative for toxin production and thus correspond to non-toxigenic toxin-gene bearing (NTTB) isolates. Six of the NTTB isolates had a stop codon within the *tox* gene sequence (**Table S1; Table S2**). However, for the 11 remaining strains, we found no explanation for the observed lack of toxin production.

Upon biotyping, 154 (62.3%) isolates belonged to biovar Mitis, 87 to biovar Gravis (35.2%) and 6 (2.4%) to biovar Belfanti (**Table S1**). Biovar proportions were similar among the three datasets. Mitis isolates were more frequently *tox*-positive than Gravis isolates (56/154 *versus* 18/87, chi-squared test, *p*-value 0.01; **Figure S1**). Among *tox*-positive isolates, NTTB were more frequent among Mitis isolates (13/56, 23.2%) than among Gravis isolates (1/18, 5.6%) although this difference was not statistically significant (p-value 0.09). Three out of four *tox*-positive Belfanti isolates were NTTB.

### Phylogenetic structure of *C. diphtheriae* and distribution of the toxin gene

To infer a phylogenetic tree, we first aimed to detect and remove homologous recombination events among *C. diphtheriae* genomic sequences. ClonalFrameML inferred a relative rate of recombination to mutation (R/theta) of 0.86, with an average length of recombination segments (delta) of 287 bp. The mean genetic distance between donor and recipient of recombination (nu) was 0.02 substitutions per nucleotide position, resulting in a relative impact of recombination to mutation (r/m=R/theta×delta×nu) of 5.01.

The recombination-corrected phylogeny (**Figure 2; Figure S2)** was star-like, with a multitude of sublineages branching off deeply. The deepest branching sublineages corresponded to two ribotype reference strains of biovar Mitis: CIP107521 (ribotype Dagestan) and CIP107534 (ribotype Kaliningrad). Remarkably, isolates of biovars Mitis and Gravis were mostly distributed in two distinct branches of the tree. We therefore named the two major branches, lineage Mitis (156 strains, of which 86% were of biovar Mitis) and lineage Gravis (91 strains, of which 77% were of biovar Gravis). The Gravis lineage branched off from within the Mitis lineage (**Figure 2**). Reference strains PW8 and NCTC11297^⊤^ belonged to the Mitis lineage, whereas NCTC13129 (from the ex-Soviet Union 1990’s outbreak) and NCTC10648 belonged to the Gravis lineage. The Belfanti isolates were scattered in three distinct sublineages within the Mitis lineage and one within the Gravis lineage.

**Figure 2.**
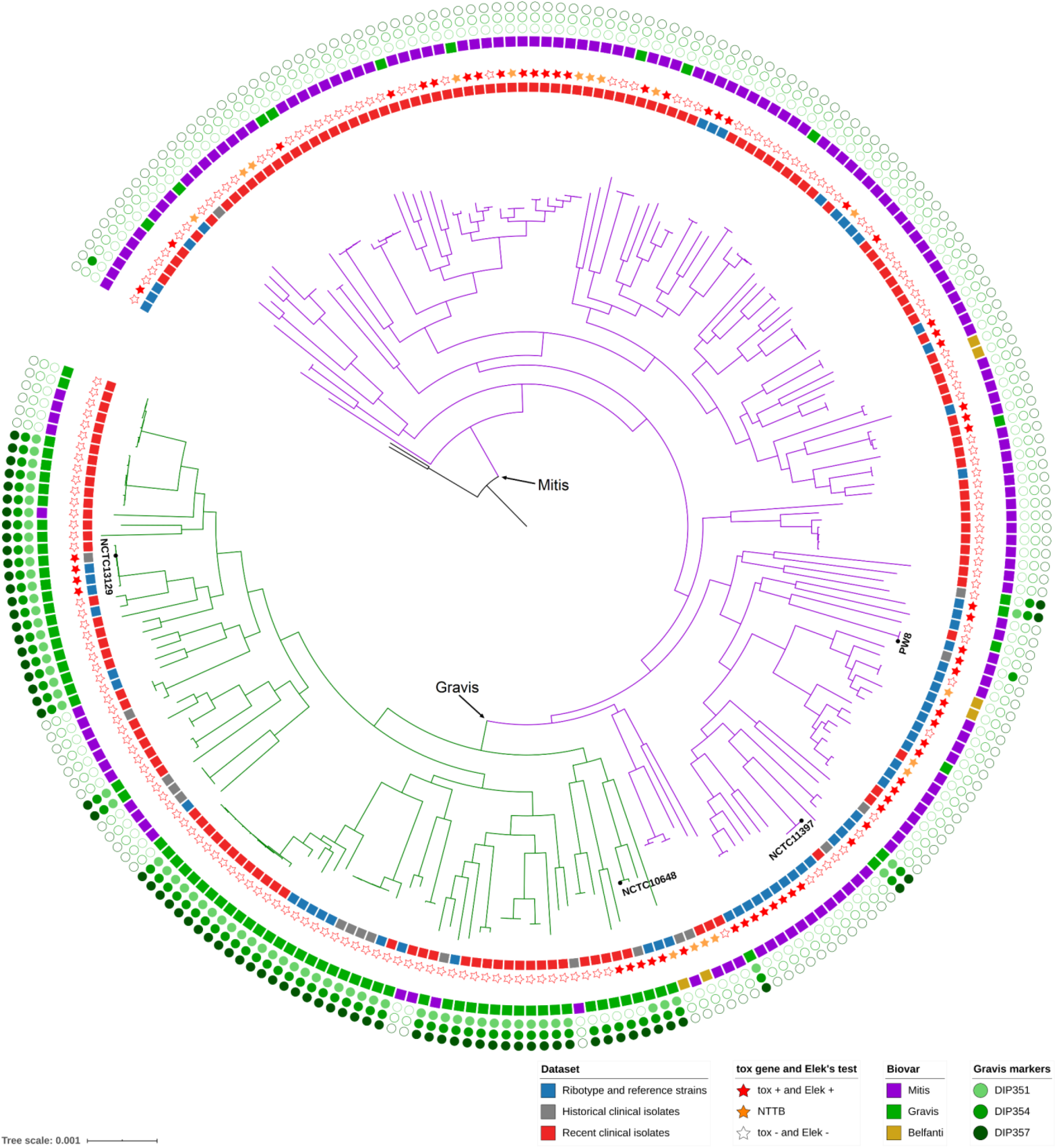
Phylogenetic tree of *C. diphtheriae*. The tree was obtained using ClonalFrameML and was rooted using *C. rouxii* and *C. belfantii* isolates (not shown). Main lineages Mitis and Gravis are labeled and their branches are drawn using purple and green, respectively. The first (internal) circle around the tree corresponds to the three strain subsets (red: recent clinical isolates; blue: reference strains; grey: older clinical isolates). The second circle (stars) gives the toxigenic status. The third circle corresponds to biovars Mitis (purple), Gravis (green) and Belfanti (yellow). The next three circles indicate the presence of the spuA-associated gene cluster; DIP357 = *spuA* gene. The positions of reference strains PW8, NCTC13129 and NCTC10648 are indicated. The scale bar give the number of nucleotide substitutions per site.

The isolates carrying the *tox* gene belonged mostly to the Mitis lineage (68 of 78, 87.2%), in which they were distributed in multiple sublineages. In the Mitis lineage, 69 (44.2%) were *tox*-positive. In contrast, within the Gravis lineage, only 10 (11%) isolates were *tox*-positive, and they corresponded to the earliest-branching Gravis sublineages with only one exception. Interestingly, this exception corresponded to the large ex-Soviet Union outbreak in the 1990s (**Figure 2**). This phylogenetic pattern is consistent with an evolutionary scenario where Mitis is the ancestral biovar of *C. diphtheriae* and where Gravis evolved from the Mitis lineage as an initially *tox*-positive sublineage, with subsequent loss of the toxin gene. In this scenario, the ex-Soviet Union outbreak sublineage would have re-acquired the *tox* gene. All NTTB isolates belonged to the Mitis lineage except strain CIPA99 (ribotype Rhone, biovar Belfanti; **Figure 2**), and they were distributed in multiple sublineages, showing convergent evolution towards the loss of toxin production.

### Genetic events linked to biovar status

Biovar Mitis and Gravis are distinguished by the ability to utilize glycogen (positive in Gravis, negative in Mitis). The *spuA* gene, which codes for a putative alpha-1,6-glycosidase, was reported as being specific for biovar Gravis isolates ^41^. Our genome-wide association study (GWAS) of accessory genes with the biovar phenotype revealed a strong association of a cluster of genes that includes *spuA* (DIP357; **Figure S3**) with biovar Gravis isolates. This association was stronger within the Gravis lineage; in contrast within the Mitis lineage, few of the biovar Gravis isolates possessed *spuA* (**Figure 2**). GWAS analysis of core SNPs further demonstrated that a SNP (at position 324,487, **Figure S3**) downstream of the *spuA* cluster insertion point was also associated with biovar, suggesting homologous recombination among core genes as a mechanism for the *spuA* cluster insertion event.

The nitrate reductase activity differentiates Mitis and Gravis isolates, which are positive, from Belfanti isolates, which are nitrate-negative. We found that the nitrate reduction *narKGHJI* gene cluster ^41^ was disrupted in three of the six isolates assigned to the biovar Belfanti: strains FRC0480 and FRC0481 had a G to A mutation at position 675 of the *narG* gene, leading to a stop codon; whereas in strain CIPA99, approximately 100 nucleotides were inserted at position 446 in *narG*. No molecular explanation was found for the lack of nitrate reductase ability of the three other Belfanti strains when scrutinizing the *narKGHJI* gene cluster and adjacent molybdenum cofactor biosynthesis genes ^42^.

### Antimicrobial susceptibility variation

Susceptibility to 19 antimicrobial agents was determined for the 247 clinical isolates and reference strains (**Table S3**). For each agent, the distribution of zone diameter (ZD) values (**Figure 3**) revealed a predominant mode located towards the right end of the distribution. This mode likely corresponds to the natural susceptibility distribution within the *C. diphtheriae* population and was used to define tentative epidemiological cutoffs (ECOFF, also called ecological cutoff ^43^). The proposed ECOFFs and their comparison with clinical breakpoints are presented in **Table S4.** For each antimicrobial agent except cefotaxime, this approach led to the identification of outsider strains with potentially acquired resistance (**Figure 3**).

**Figure 3.**
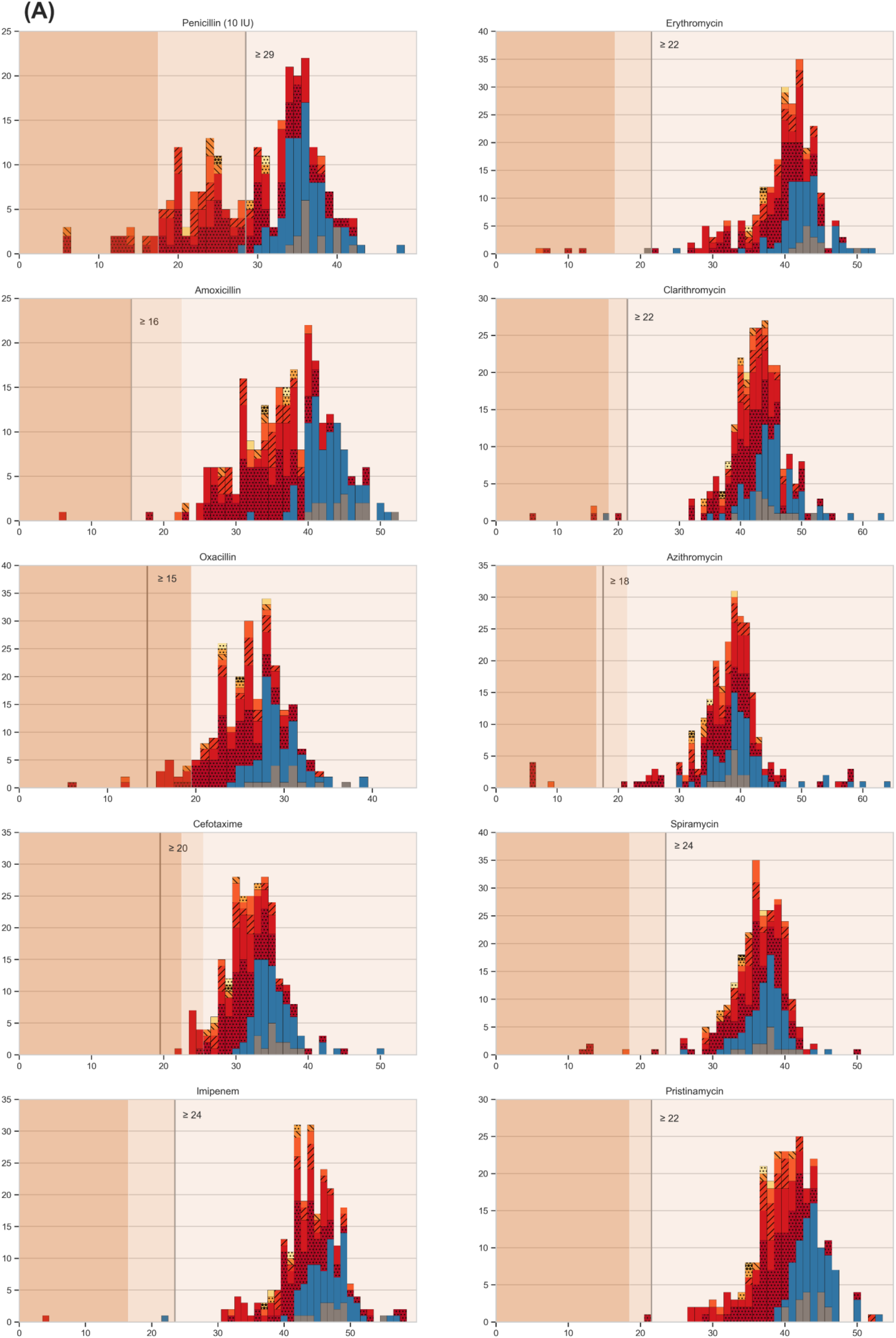
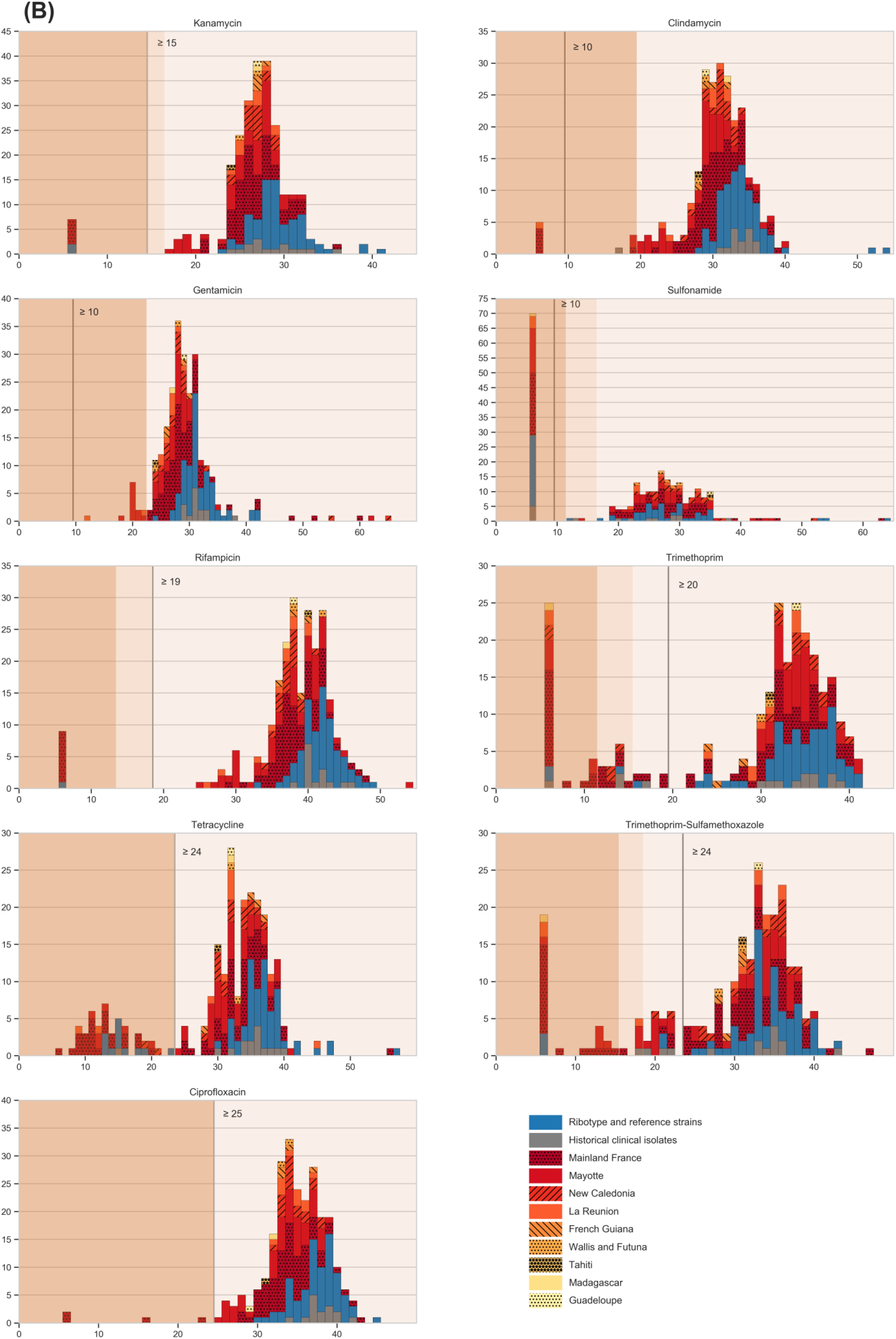
The distributions of zone diameter values for 19 antimicrobial agents. **A**: beta-lactams, macrolides and pristinamycin. **B**: other agents. X-axis: diameter in mm; Y-axis: number of strains. Colors inside the bars represent subset and geographic origins as in Figure 1 (see key on panel B). The three background colors represent the categorical interpretations according to EUCAST: resistant (salmon, left), intermediate (lighter salmon, middle) and susceptible (pale beige, right). The grey vertical bar corresponds to the proposed tentative ecological cutoff.

Penicillin was exceptional in that the predominant susceptible mode (centered around 36 mm) was less neatly defined, due to partial overlap with a second mode of smaller diameter values centered around 24 mm. This second mode corresponds mostly to the ‘intermediate’ interpretative category (18<=ZD<29 mm) but also overlaps with the ‘resistant’ category (< 18 mm). The distribution of ZD values for tetracycline also showed a clear second mode. For multiple other agents (amoxicillin, oxacillin, imipenem, kanamycin, rifampicin, ciprofloxacin, clindamycin and more evidently sulfonamide, trimethoprim and the trimethoprim-sulfamethoxazole combination), outsider strains had the minimal diameter (6 mm, corresponding to growth at the disk contact). For trimethoprim, we observed both a mode centered around 14 mm and a group of even more resistant outliers with growth at disk contact.

Antimicrobial resistance levels were similarly distributed between *tox*-positive and *tox*-negative isolates (**Figure S4**) as well as between the two main phylogenetic lineages or biovars (**Figure S5; Table S1**).

Resistance rates were 17.2%, 2.5% and 2.5% for penicillin, amoxicillin and erythromycin, respectively, among the prospectively collected 2008-2017 clinical isolates (**Figure 4**). Reference strains were generally susceptible to most agents, including penicillin, but were partially resistant to tetracycline (18%) and sulfonamide (35%). The resistance profiles distribution showed that approximately half (121/247) of the strains had a fully susceptible phenotype, whereas four isolates were multidrug resistant (> 8 agents; **Figure 4 inset**). Notably, these four isolates were resistant at the same time to penicillin and macrolides, and two of them (FRC0402 and FRC0466) additionally had a reduced susceptibility to amoxicillin. Two multidrug resistant isolates were collected from a foot arch wound and in respiratory carriage in the same patient (French mainland, with recent travel from New Caledonia). The two others came from a patient living in La Réunion Island (FRC0402) and from a patient living in Paris, who had recently traveled to Tunisia (FRC0466). Isolates from La Réunion Island and mainland France showed resistance to multiple antimicrobial agents more often than isolates from other geographic origins (**Figure 1A inset**).

**Figure 4.**
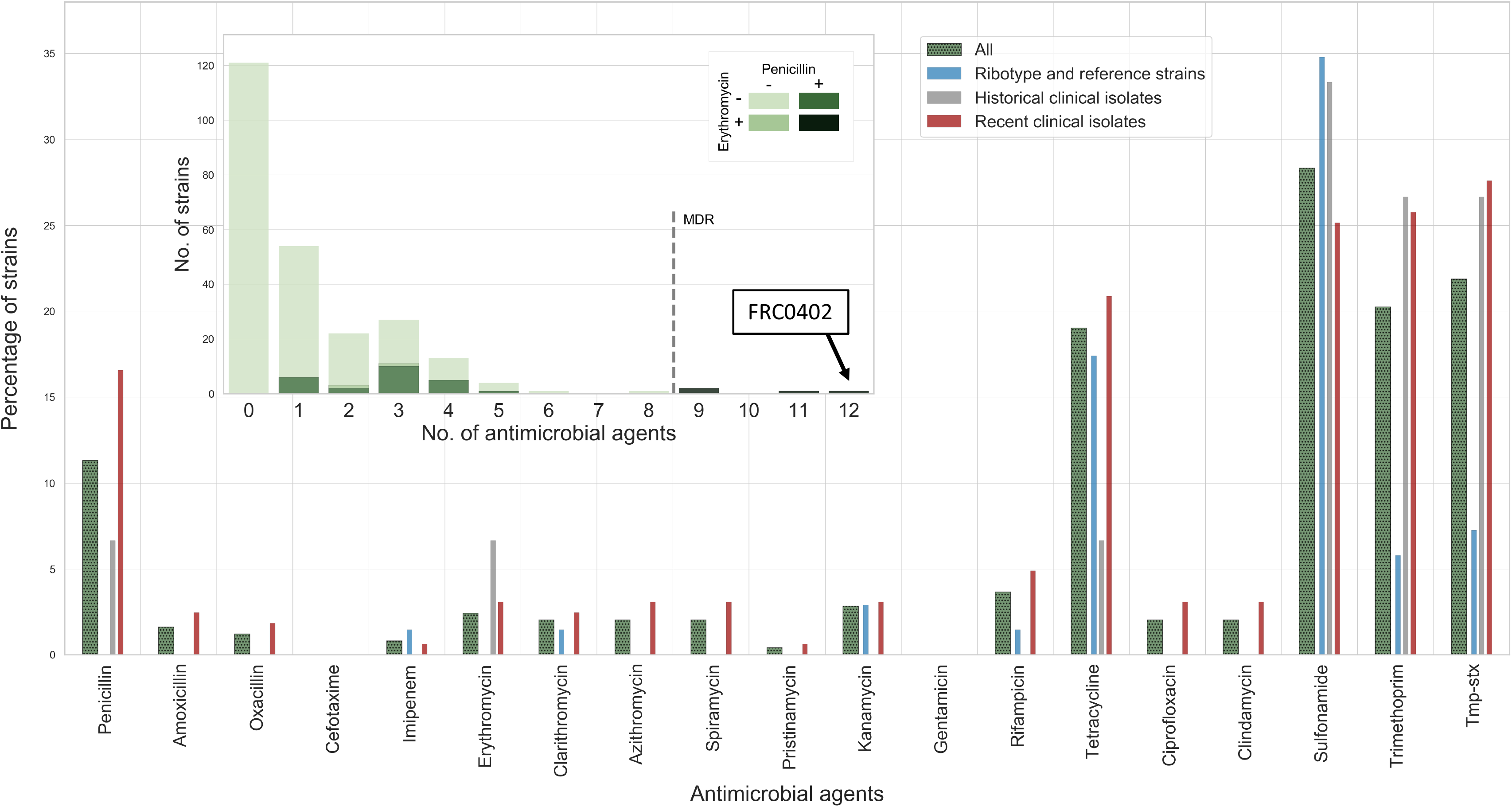
Proportions of resistant strains by antimicrobial agent and multidrug resistance phenotypes. Interpretation of zone diameter values was performed according to proposed ecological cutoffs. The main panel shows the percentage of strains resistant to each agent. Tmp-stx: trimethoprim-sulfamethoxazole. The four bars for each agent correspond to the entire dataset (all, shaded green) or the three subsets (see key). The inset shows the number of strains resistant to a given number of antimicrobials. Penicillin and/or erythromycin resistant strains are colored with darker grey (see key). The vertical bar indicates the definition of multidrug resistant isolates (> 8 agents). FRC0402, the most multidrug resistant isolate with resistance to 12 agents, is highlighted.

### Genomic associations with antimicrobial resistance phenotypes

We first searched for the presence in the genomic sequences, of previously described antimicrobial resistance genes (ARGs). This approach led to detection of 12 ARGs (**Table S5**). We identified three tetracycline resistance genes (*tetW, tet33* and *tetO*), four aminoglycoside resistance genes [aph(3’)-Ia, aph(3’’)-Ib, aph(6)-Id, and aadA1 = ant(3’)-Ia], and also *ermX*, *dfrA16, dfrA15b, dfrA1* and *sul1* genes. We observed a strong correlation between the presence of ARGs and the expected resistance phenotypes (**Figure S6**), particularly for *ermX* (macrolide resistance), *sul1* (sulfonamide resistance) and *aph(3’)-Ia* (kanamycin resistance). In strains FRC0137 and FRC0375, this latter gene was linked to *aph(3’)-Ib* (strA), *aph(6)-Id* (strB) and *ermX* on a Tn5432-like genomic region with an IS1249 insertion sequence ^44^. The phylogenetic distribution of ARGs (**Figure 5**) revealed their presence in multiple unrelated sublineages, consistent with independent acquisitions by horizontal gene transfer. Gene *ermX* was present either in proximity to gene *pbp2m* (see below) or in a fragmented insertion-sequence rich accessory region. Gene *dfrA16* was associated with *sul1* on a reported ^38^ class 1 integron (see below, resistance plasmid section). Tetracycline resistance was associated either with the ribosomal protection protein genes tet(O) or tet(W), or with the efflux pump gene tet33. These three genes were present in distinct strain subsets and appear to contribute independently to tetracycline resistance in *C. diphtheriae* (**Figure 5**); they were mostly associated with insertion sequences but not with other ARGS.

**Figure 5.**
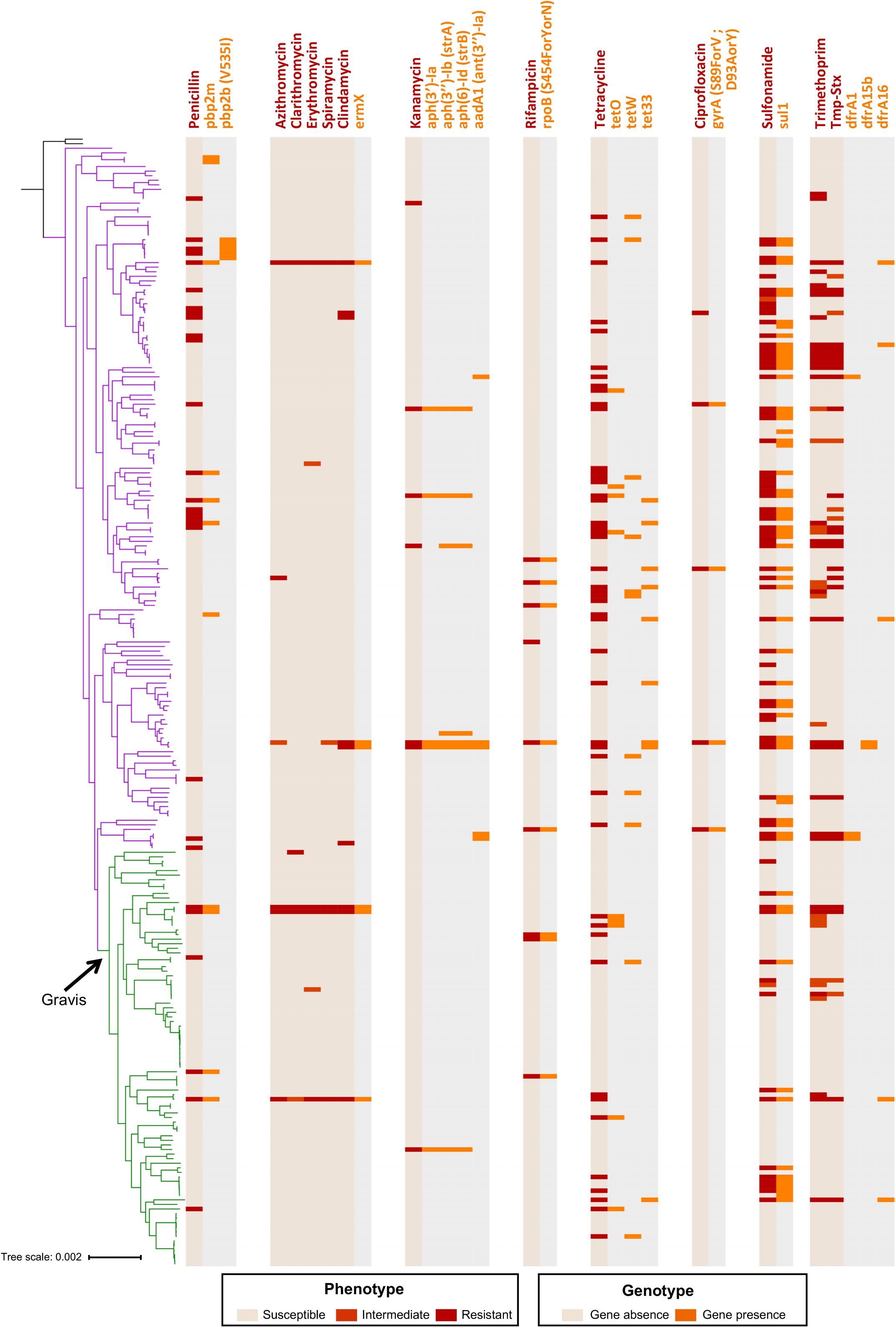
Phylogenetic distribution of antimicrobial resistance phenotypes and genes. The phylogenetic tree is the same as in Figure 1; the Mitis branch is in purple, the Gravis branch in green. To the right of the tree, each bloc indicates first, the phenotype (resistant: red; see key) and relevant corresponding genotypes (orange: gene or mutation presence). The last bloc shows resistance genes linked to chloramphenicol, which was not tested phenotypically.

Next, in order to identify novel genetic determinants potentially associated with antimicrobial resistance in *C. diphtheriae*, a GWAS approach was followed, based on either core genome SNPs or accessory gene presence/absence. SNPs that were strongly associated with ciprofloxacin, trimethoprim and rifampicin resistance were identified within genes for gyrase subunit A, dihydrofolate reductase and RNA polymerase subunit B, respectively (**Table S5; Figure S7**), consistent with known mechanisms and validating our approach. SNPs were also found to be associated with penicillin, kanamycin and tetracycline resistance (**Table S5**), but in these cases the functional attribution is undefined. No association was found for penicillin-resistance within the core PBP genes using the genome-wide approach. However, using a concatenation of the amino-acid sequences of the seven identified putative PBPs of *C. diphtheriae*, we identified amino-acid positions that were statistically associated with penicillin resistance (**Figure S8**). The identified positions were mapped onto the predicted functional domains of the different PBPs (**Figure S9**) revealing several mutation hotspots. A number of significant SNPs were found within the transpeptidase (TP) domains of the different PBPs, but none of the mutations mapped to the conserved transpeptidation motif. Other mutations were found outside the TP domains, for instance in the transglycosylase and PASTA domains of PBP1b or in the dimerization domain of PBP2b.

GWAS analysis of accessory genes demonstrated significant associations with phenotypic resistance. Associated genes included those mentioned above for erythromycin, tetracycline, kanamycin, sulfonamide and trimethoprim (**Table S5**). In addition, an accessory penicillin-binding protein gene (which we name *pbp2m*) was strongly associated with penicillin resistance. This gene is described in more detail below.

### Discovery of a novel penicillin-binding protein (PBP2m) associated with penicillin resistance

The novel PBP gene *pbp2m* was observed in 11 isolates, 8 of which were penicillin resistant with minimum inhibitory concentrations (MIC) ranging from 0.19 to 1.5 mg/L. Two of these isolates were also resistant to amoxicillin and one was in addition resistant to oxacillin (**Table S4**). The phylogenetic distribution of *pbp2m*-positive strains was compatible with multiple independent acquisitions of the gene through horizontal gene transfer (**Figure 5**). Sequence analysis showed that the newly identified PBP2m is almost identical to PBP2c from *C. jeikeium*, a class B PBP with an N-terminal signal peptide followed by a lipobox domain and the C-terminal transpeptidase domain (**Figure S9**). The *C. jeikeium* PBP2c is a low affinity PBP and was associated with beta-lactam resistance in *C. jeikeium* ^45^.

To demonstrate the role of PBP2m in penicillin resistance, its gene was PCR amplified from FRC0402 and cloned into the pTGR5 plasmid (**Figure S10**). Transformation of the plasmid into *C. glutamicum* strain ATCC 13032 raised the MIC for penicillin from 0.125 to 1.5 mg/L, and the MICs of the other beta-lactams amoxicillin, cefotaxime and oxacillin also increased importantly **(Figure 6; Table S6**). In contrast, MICs of non-beta-lactam agents were not changed. Imipenem was less effective against the transformant based on disk diffusion but not based on E-test. Transformation with the empty plasmid used as control did not affect the MIC of any agent. These results show that PBP2m confers resistance to a broad range of beta-lactams.

**Figure 6.**
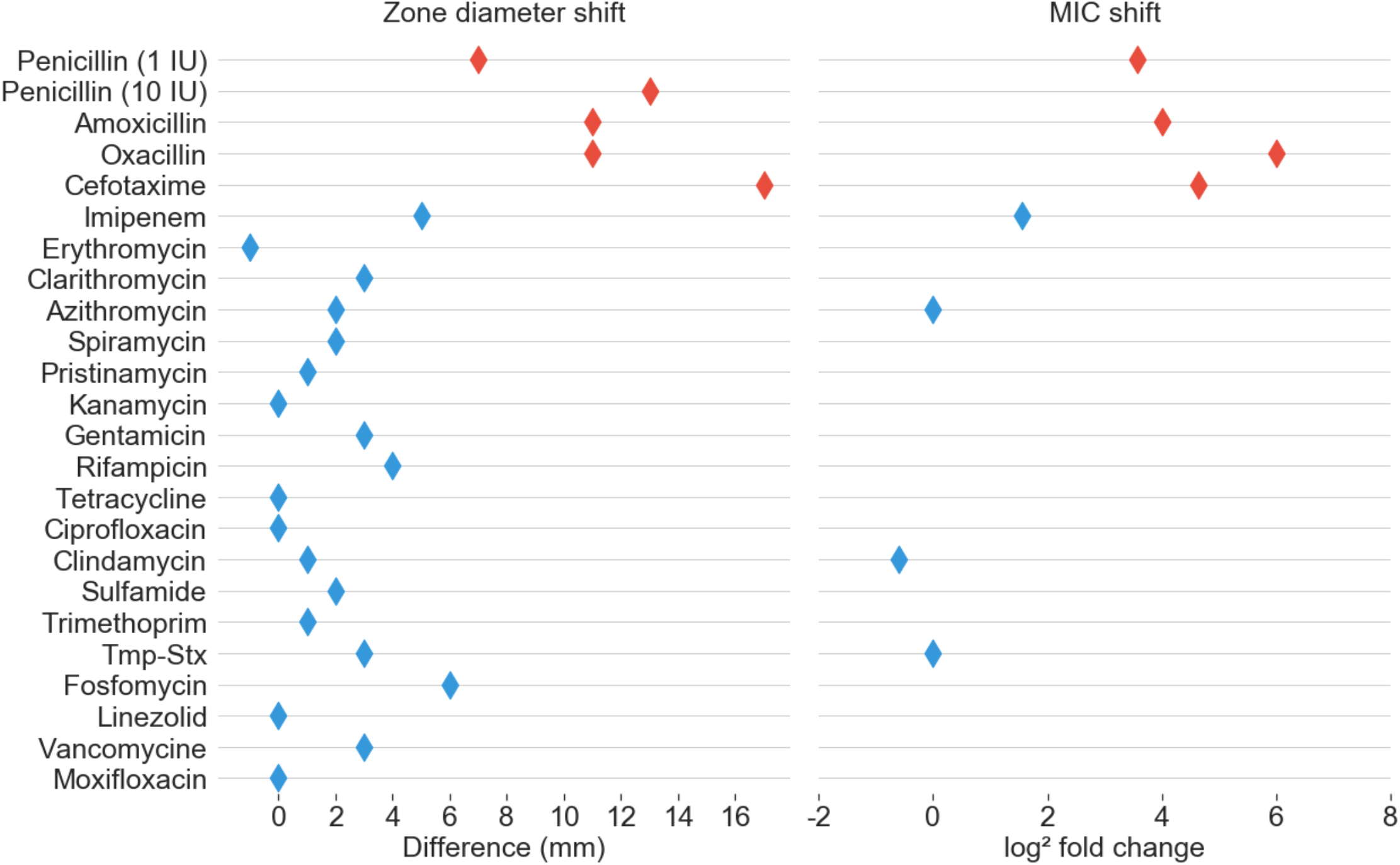
Phenotypic effect of *pbp2m* expression. Compared susceptibility phenotypes for *C. glutamicum* transformants with plasmid pTGR5 containing, or not, the *pbp2m* gene. Left, shift in zone diameter size; right, shift in the minimum inhibitory concentration (MIC). Diamonds are positioned on the scales, at positions corresponding to the difference of zone diameters (without *pbp2m* – with *pbp2m*) or the log2 of MIC ratios (with *pbp2m*/without *pbp2m*). Red, penicillins or cephalosporins; blue, other agents. Tmp-Stx: trimethoprim-sulfamethoxazole.

### Discovery of a multidrug resistant conjugative plasmid carrying the gene *pbp2m*

Strain FRC0402, a *tox*-negative isolate from La Réunion Island, stood out as being resistant to 12 agents (**Figure 4 inset**). In addition to *pbp2m*, this isolate carried genes *sul1, ermX* and *dfrA16* and a *tetA* family *tet(Z)*-like (71%) tetracycline efflux gene. To define the genomic context of resistance genes, a complete genome sequence was obtained. The assembly revealed a chromosome of 2,397,465 bp and a circular plasmid of 73,763 bp (**Figure 7**), which we propose to name pLRPD (for large resistance plasmid of *C. diphtheriae*).

**Figure 7.**
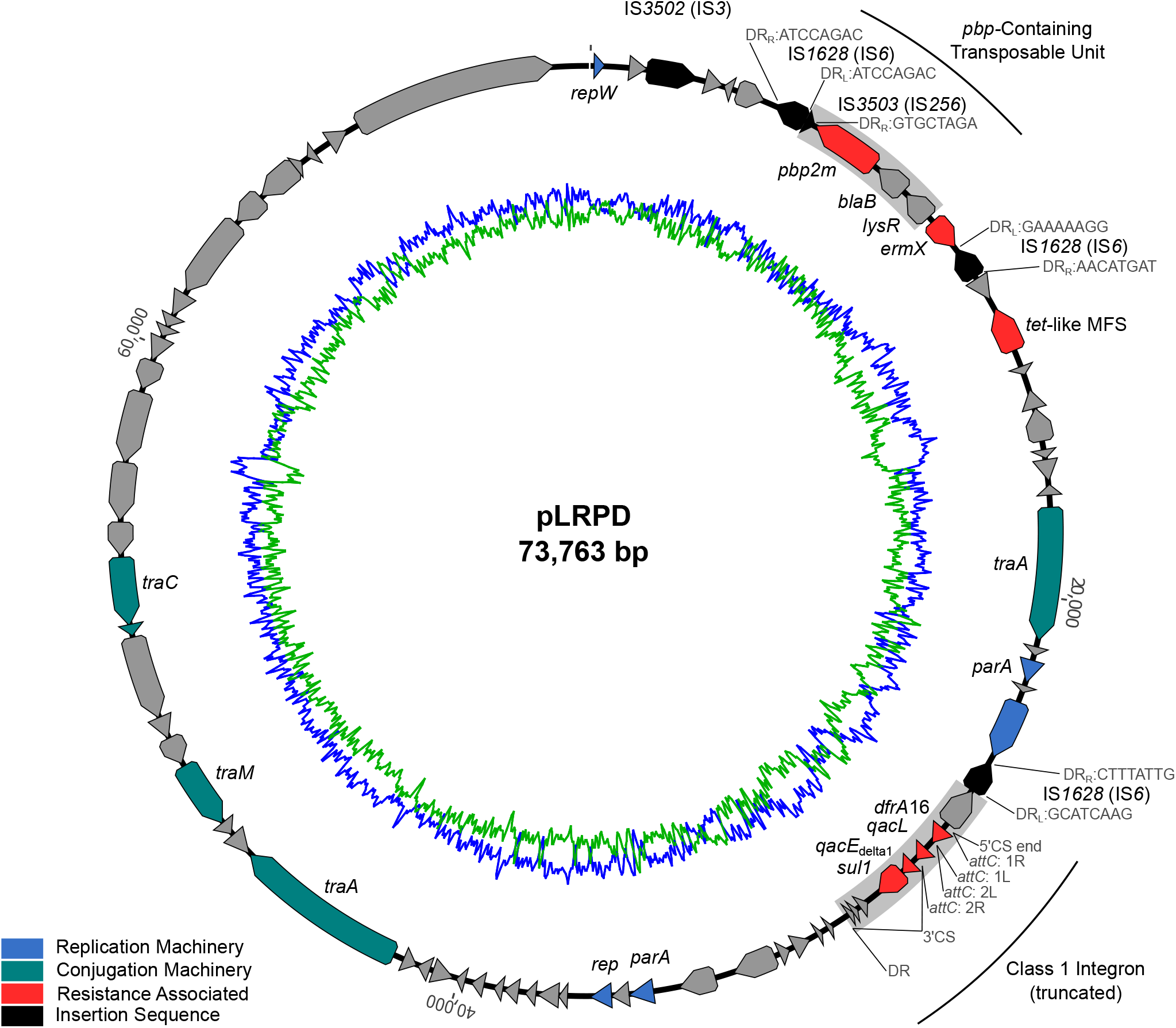
Map of plasmid pLRPD from isolate FRC0402. FRC0402 Predicted coding sequences are portrayed by arrows and coloured based on the predicted gene function (refer to key). Inner blue circle, G+C% content; inner green circle, A+T% content. IS3502 annotation is putative; IS3503 is truncated.

The *pbp2m* gene was located on the large plasmid in a region comprising three other genes, a *blaB* beta-lactamase family gene, a LysR family regulator gene and the *ermX* gene, flanked by two insertion sequences (IS*1628*) of the IS*6* family (**Figure 7**). With disparate direct repeat sequences, it remains unclear if this region represents a single transposable unit or a mosaic of gene acquisition events in pLRPD. A nearly identical PBP was observed in 78% of publicly available *C. jekeium* genomes, in 57% of *C. striatum* genomes and in multiple other *Corynebacterium* genomes (**Table S7**). However, the genetic context of PBP2m was highly variable in *C. diphtheriae* and among other *Corynebacterium* species (**Figure S11**). A putative transposable PBP-containing unit (PCU) comprising genes *pbp2m*, *blaB* and *lysR*, commonly flanked by IS*3503* (IS*256* family) with a fragment identified in pLRPD (**Figure 7**), appeared to be highly conserved and was associated variably with *ermX* in *C. diphtheriae* and with a helicase in *C. diphtheriae* and other *Corynebacterium* species. The PCU was sometimes found in 2 or 3 tandem copies and was chromosomally located in most genomes.

Further elements carried by pLRPD included an integron carrying genes *dfrA16*, *qacL*, *sul1*, as well as elements of a putative conjugation apparatus gene cluster (**Figure 7**). Our conjugation experiments aiming to demonstrate the transfer of pLRPD into recipient *C. diphtheriae* isolates failed. This plasmid was not found in other *C. diphtheriae* strains.

## DISCUSSION

Strains of *C. diphtheriae* that are resistant to antimicrobial therapy may compromise the management of diphtheria cases and the control of pathogen transmission. Here we aimed to define the genomic determinants of resistance to penicillin and other antimicrobial agents in *C. diphtheriae*, and to analyze the relationships of resistance with diphtheria toxin production, biochemical variants and phylogenetic sublineages. To this aim, we characterized phenotypically and genotypically, a large sample of *C. diphtheriae* isolates from diverse geographic and temporal origins. We confirmed that the species is made of multiple phylogenetic sublineages^9,13,46^ and showed that homologous recombination contributes five times more to their diversification than mutation, consistent with previous evidence of recombination in *C. diphtheriae* populations ^12,13^.

Historically, *C. diphtheriae* isolates have been classified into three main biovars, but the links between biovars and phylogenetic structure have remained obscure. Whereas previous work concluded on the absence of an association ^11^, our phylogenetic analyses reveal that Gravis and Mitis, the two main biovars of *C. diphtheriae*, are associated strongly with two phylogenetic lineages. Lineage Gravis appears to have acquired ancestrally a gene cluster comprising the extracellular glycogen debranching enzyme gene *spuA* ^41^. Although the Gravis phenotype is largely associated with *spuA* within lineage Gravis, other genomic determinants of glycogen utilization remain to be discovered within the Mitis lineage. Whereas most of our biovar Belfanti isolates were excluded from this work because they belonged to *C. belfantii* or *C. rouxii*, a few Belfanti isolates did belong to *C. diphtheriae*. Our results show that biotyping is subject to parallel evolution and has limited epidemiological typing value.

The most important factor of *C. diphtheriae* pathogenicity is the diphtheria toxin. Despite early realization that it is encoded on a prophage ^5^, few studies have investigated the phylogenetic distribution of the *tox* gene in *C. diphtheriae* ^8,9^. Here, we show that *tox*-positive strains mainly belong to the Mitis lineage and to early-diverging branches of the Gravis lineage. The distribution of *tox*-positive isolates into multiple Mitis sublineages is strongly indicative of independent acquisitions of the toxin gene. Alternately, this pattern might result from initial acquisition of the *tox* gene, followed by secondary loss in multiple sublineages. The phylogenetic pattern is also consistent with an ancestral presence of the *tox*-bearing phage in the Gravis lineage, with subsequent loss of the *tox* gene in the branch leading to the ancestor of most Gravis isolates. Future work should investigate the dynamics of the lysogenic corynephages and molecular determinants of their sublineage distribution. One important open question is the likelihood of *tox*-negative strains acquiring the *tox* gene during colonization, infection or short-term epidemiological timeframes ^7^.

Remarkably, except for early-diverging sublineages, only one Gravis sublineage was found to carry the *tox* gene. This sublineage happens to correspond to the largest outbreak in recent times, which occurred in Newly Independent States of the ex-Soviet Union in the 1990s ^13,14,39^. This sublineage, which comprises the ST8 reference strain NCTC13129, is genetically distant from other *tox*-positive lineages, which belong to the Mitis lineage. Hence, its antigenic structures or other pathogenicity properties may have diverged from those of more common *tox*-positive isolates, which might have contributed to its exceptional transmission in the 1990s, in addition to the decline in vaccine coverage ^14^. Of note, biovar Gravis was named to reflect a perceived higher severity of infection compared to diphtheria cases caused by biovar Mitis isolates ^47,48^. Recently it was shown that most diphtheria vaccines contain, besides the anatoxin, multiple other *C. diphtheriae* immunogens ^22^. The impact of vaccination on the evolution of *C. diphtheriae* populations, and possible variations of cross-protection as a function of strain diversity, are currently undefined. This work provides a framework onto which future studies can build to address this important question.

Although antimicrobial resistant *C. diphtheriae* strains have been reported on numerous occasions ^23,32^, knowledge on antimicrobial resistance in *C. diphtheriae* is largely fragmented and suffers from lack of harmonization. Breakpoints used to define resistance vary according to world region and have changed over time within single countries ^49^. The lack of consensus on the definition of resistance restricts our ability to define the magnitude of the problem and its global significance.

We aimed to define biologically meaningful cutoffs ^43^ based on susceptibility phenotypes distributions, taking advantage of our large and diverse sample. Our data allowed us to propose tentative ecological cutoffs for *C. diphtheriae*. Clearly, this approach should in the future be extended to MIC values and should use larger and more diverse strain collections. Nevertheless, our analyses suggest that non-susceptibility to at least one antimicrobial agent was acquired by half of *C. diphtheriae* strains, regardless of lineage, biovar or toxigenic status. This study further suggests that acquired resistance to penicillin, the first line therapy against diphtheria, is far from being rare, affecting >15% of *C. diphtheriae* isolates collected in the last decade in France and its overseas territories. The high prevalence of resistance to penicillin, tetracycline and trimethoprim/sulfamethoxazole found here are consistent with susceptibility surveys of recent *C. diphtheriae* isolates in Algeria ^29^, Indonesia ^31,50^ and India ^51^. Many high-income countries such as France have chosen to use amoxicillin as the first choice for antibiotic therapy ^52^, as this molecule remains highly active. Still, widespread penicillin resistance is concerning, since diphtheria mainly occurs in resource-poor settings where penicillin G is largely used. In contrast, resistance to erythromycin and other macrolides remains rare. Our results call for concerted research into the magnitude of the antimicrobial resistance threat in *C. diphtheriae*.

Knowledge of the genetic mechanisms of antimicrobial resistance is critical for defining appropriate treatments, refining diagnostics and conducting epidemiological studies of antimicrobial resistance. Resistance genes to several antimicrobial classes have been described in *C. diphtheriae* ^37,38,53^, while additional genes described in other *Corynebacterium* species ^54^ might also be present in *C. diphtheriae*. Here, we defined the prevalence and phylogenetic distribution of previously reported and newly identified resistance determinants in *C. diphtheriae*. We demonstrate the co-occurrence of resistance phenotypes and genes, suggesting a causative link in multiple instances. We further show that resistance genes have been acquired independently in multiple sublineages, demonstrating a dynamic resistome in *C. diphtheriae*. In addition, we demonstrate an association between alterations in chromosomally encoded targets and phenotypic resistance for quinolone, trimethoprim and rifampicin. Fluoroquinolone resistance was previously linked to mutations in the *gyrA* gene in *C. amycolatum* ^55^, *C. striatum* ^56^ and *C. belfantii* ^57^ but seemingly never for *C. diphtheriae*. Finally, we demonstrate the co-occurrence within some strains, of multiple resistance determinants and uncover a previously undescribed large resistance plasmid in *C. diphtheriae*. The mechanism of genetic transfer of this plasmid remain to be investigated. This work provides a first overview of the *C. diphtheriae* resistome and will facilitate further studies into the evolutionary emergence of multiresistant *C. diphtheriae* strains.

Mechanisms of penicillin resistance in *C. diphtheriae* have never been described, to our knowledge. Here we discovered an accessory PBP (PBP2m), which was experimentally shown to confer resistance to penicillin and other beta-lactam antimicrobial agents. Its distribution in multiple sublineages, and its presence in other *Corynebacterium* species, clearly demonstrates its horizontal transfer, and we revealed a multiplicity of genomic contexts in which it is found within *Corynebacterium*. PBP2m is a putative low affinity PBP, which would explain why it is less affected by beta-lactam antibiotics. Further studies on the expression, antimicrobial resistance spectrum and mechanism of action of PBP2m are warranted.

The seven chromosomal PBPs of *C. diphtheriae* (including the newly annotated PBP4b) were investigated to identify amino acid sequence polymorphisms associated with penicillin resistance. Although several alterations were significantly associated, none were directly implicated with the catalytic residues of the transpeptidase or transglycosylase domains. The association with resistance may not be directly linked to these catalytic residues but could be due to secondary sites that are thought to interfere with beta-lactam ligand binding. While biochemical and structural studies are necessary to understand how these mutations affect penicillin susceptibility, we postulate that some changes in these domains could lead to allosteric effects ultimately resulting in beta-lactam resistance, as described for *Staphylococcus aureus* PBP2a ^58,59^ or *Streptococcus pneumoniae* PBP2x ^60^. Other SNPs might simply have been hitchhiking due to their physical linkage with functionally important SNPs ^61^.

## Conclusion

As a result of vaccination and antitoxin therapy, diphtheria has fallen from a main killer of young children to a largely controlled disease. However, in recent years vaccination rates have dropped in several settings afflicted by conflicts or economic crises, and the lack of availability of diphtheria antitoxin is becoming critical. Antimicrobial therapy is an increasingly important component of diphtheria control, but its efficacy is jeopardized by emerging resistance. Here we contributed to define the magnitude of this issue and provide novel insights into its genomic underpinnings. We also provide fresh views on the population structure of the *C. diphtheriae* species, and associations between traditional biochemical characterization of strains into biovars, the distribution of toxigenic isolates, and the population dynamics of antimicrobial resistance within *C. diphtheriae*.

## METHODS

### *Corynebacterium diphtheriae* isolates and strains

A collection of 247 *C. diphtheriae* isolates were included (**Table S1**), corresponding to three subsets. First, we included 163 clinical isolates (Recent clinical isolates subset, **Table S1**) collected prospectively between 2008 and 2017 by the French National Reference Center for Corynebacteria of the *Corynebacterium diphtheriae* complex (NRC-CCD). These isolates represented all isolates received at the NRC-CCD that corresponded to the *C. diphtheriae* species (*C. belfantii, C. ulcerans, C. pseudotuberculosis* or other corynebacteria were excluded). They were collected from cutaneous (n = 136), respiratory (n = 23) and other infections (bones, blood; n = 4). Of these, 74 were from Mainland France and 89 from French overseas territories, including Mayotte (n=50), New Caledonia (n=19), La Réunion island (n=11), French Guiana (n=4), French Polynesia (n=3) and Guadeloupe (n=1); one isolate received from Institut Pasteur in Madagascar was also included (**Figure 1**). Four isolates from New Caledonia collected between 2002 and 2006 (02-0322, 02-0338, 03-1641 and 06-1569) were included in a previous study ^62^; the trimethoprim and sulfamethoxazole-resistant isolate FRC0024 was previously shown to harbor an integron with gene *drfA16* ^38^.

Second, we included 15 clinical isolates collected in France between 1981 and 1991, 11 of which had been deposited in the Collection de l’Institut Pasteur (CIP; Historical clinical isolates subset in **Table S1**).

Third, to increase the genetic diversity and geographic range of the sample, the 65 available reference strains of ribotypes that belong to *C. diphtheriae* were included ^40^. These reference strains represent an international collection of isolates collected over several decades and originating from multiple world regions including the Americas, Europe, Asia, Africa, and Oceania. Our subcultures of these strains were controlled for *tox* gene presence, toxin production and biovar, leading to modifications of published characteristics in some instances (**Table S1**; **Figure 1**). Finally, four reference strains were included: strain NCTC13129, which is used as genomic sequence reference ^39^; strain NCTC10648, which is used as the *tox*-positive and toxinogenic reference strain in PCR and Elek tests, respectively; strain NCTC11397^⊤^, which is the taxonomic type strain of the *C. diphtheriae* species; and the vaccine production strain PW8, which corresponds to CIP A102 ^63^. This third subset is referred to as “Ribotype and reference strains” subset (**Table S1**).

### Bacterial cultures, identification and biovar

Bacteria were cultivated on Trypto-Casein-Soy (TCS) agar during 24 h at 35-37°C. Bacterial identification was performed at the NRC-CCD as described previously ^64^ by multiplex polymerase chain reaction (PCR) combining a *dtxR* gene fragment specific for *C. diphtheriae* and a multiplex PCR that targets a fragment of the *pld* gene specific for *C. pseudotuberculosis*, the gene *rpoB* (amplified in all species of the *C. diphtheriae* complex) and a fragment of 16S rRNA gene specific for *C. pseudotuberculosis* and *C. ulcerans*. Isolates collected since 2014 were confirmed as *C. diphtheriae* by matrix-assisted laser desorption-ionization time-of-flight mass spectrometry (MALDI-TOF MS) using Bruker technology. In order to exclude strains initially identified as *C. diphtheriae* but now classified as *C. belfantii* ^64^ or *C. rouxii* ^65^, genome-wide average nucleotide identity (ANI) was used as described previously ^64^. Strains were characterized biochemically for pyrazinamidase, urease, nitrate reductase and for utilization of maltose and trehalose using API Coryne strips (BioMérieux, Marcy l’Etoile, France) and the Rosco Diagnostica reagents (Eurobio, Les Ulis, France). The Hiss serum water test was used for glycogen fermentation. The biovar of isolates was determined based on the combination of nitrate reductase (positive in Mitis and Gravis, negative in Belfanti) and glycogen fermentation (positive in Gravis only). The rare biovar Intermedius was not identified, as its distinction from other biovars is based on colony morphology, which is considered subjective, or on lipophily, which was not tested.

### Determination of the presence of the *tox* gene

Determination of the diphtheria toxin gene (*tox* gene) presence was achieved by a conventional *tox* PCR assay ^66^, while its phenotypic production was assessed by the modified Elek test ^67^. We also confirmed *tox* PCR results by BLASTN (query: *tox* gene sequence from strain NCTC13129, RefSeq accession number: DIP_RS12515) analysis of the genomic assemblies.

### Antimicrobial susceptibility testing

Phenotypic susceptibility was tested for the following agents: penicillin G (10 IU), amoxicillin, oxacillin, cefotaxime, imipenem, erythromycin, azithromycin, clarithromycin, spiramycin, pristinamycin, kanamycin, gentamicin, rifampicin, tetracycline, ciprofloxacin, clindamycin, sulfonamide, trimethoprim, and trimethoprim + sulfamethoxazole. The 19 antimicrobial agents tested (Table S1) corresponded to seven classes, as described hereafter. *β*-lactams: penicillin G (PEN), amoxicillin (AMX), oxacillin (OXA), cefotaxime (CFT), imipenem (IMP); Macrolides: azithromycin (AZM), clarithromycin (CLR), erythromycin (ERT) and spiramycin (SPR); Lincosamides: clindamycin (CLD); Streptogramins: pristinamycin (PRT); Aminoglycosides: gentamicin (GEN) and kanamycin (KAN); Folate pathway inhibitors: sulfonamide (SUL), trimethoprim (TMP) and trimethoprim + sulfamethoxazole (cotrimoxazole, TMP-STX); Ansamycins: rifampicin (RIF); Tetracyclines: tetracycline (TET); and Fluoroquinolones: ciprofloxacin (CIP).

Antimicrobial susceptibility was determined using the disk diffusion method with impregnated paper disks (Bio-Rad, Marnes-la-Coquette, France) on Mueller Hinton agar plates supplemented with 5% of sheep blood and 20 mg/L β-NAD, as recommended. Minimum inhibitory concentrations (MIC) were determined using E-test strips (BioMerieux, Marcy l’Etoile, France). The control strain used is *S. pneumoniae* ATCC 49619. The zone diameter (ZD) data were interpreted into S, I and R categories in the following way. First, we used the CA-SFM/EUCAST V.1.0 (Jan 2019) document (https://www.sfm-microbiologie.org/wp-content/uploads/2019/02/CASFM2019_V1.0.pdf), which contains interpretative criteria for *Corynebacterium* spp. only for CIP, GEN, CLD, TET, RIF and TMP-STX. Second, for the other agents, we used the interpretative criteria published in Table III of the CA-SFM 2013 recommendations (https://resapath.anses.fr/resapath_uploadfiles/files/Documents/2013_CASFM.pdf). Note that for RIF, we used the 2013 breakpoints, as they fitted better with the observed distribution of ZD values. Clarithromycin breakpoints were taken from those for erythromycin, as recommended. The breakpoint for oxacillin was derived from the one used for *Staphylococcus* spp. ZD interpretation breakpoints are given in **Table S4**.

Penicillin susceptibility was initially determined using 10 UI (6 micrograms) disks (resistance breakpoint: 18 mm), but CA-SFM/EUCAST recommendations were changed in 2014 to use 1 UI disks, while the resistance breakpoint was increased from 18 to 29 mm. As all *C. diphtheriae* strains end up in the resistant category following this recommendation, E-test strips were used to define the penicillin MIC since 2014 (**Table S1**: isolates starting from FRC0259); the EUCAST breakpoint of 0.125 g/L was used as cutoff. Penicillin E-test was also performed systematically for strains tested as resistant before 2014 (using 10 UI disks), as well as for some susceptible isolates (**Table S1**).

Multidrug-resistant *C. diphtheriae* (MDR-DIP) were defined as strains resistant to more than eight of the agents tested herein, excluding intrinsic resistance to fosfomycin. Note that we used ecological cutoffs rather than currently proposed clinical breakpoints (see **Table S5** for a comparison of both types of breakpoints).

### Whole-Genome Sequencing by Illumina and Oxford Nanopore Technologies

DNA was extracted from broth cultures, by making use of DNeasy Blood & Tissue Kit (QIAGEN, Hilden, Germany). However, a lysis step was added to the extraction protocol described by the manufacturer as previously described ^68^: a 1μL loopful of bacterial colonies was emulsified in 180 μL of lysis buffer containing 20 mM Tris-HCl, pH8, 2 mM EDTA, 1.2% Triton X-100, 20 mg/mL lysozyme, in a DNase/RNase free 1.5 ml Eppendorf tube and incubated in a heating block at 37°C for 1 hour, with mixing every 20 min. After extraction, DNA concentration was measured with the Qubit 3.0 Fluorometer (Invitrogen), employing the Qubit dsDNA BR Assay Kit (Invitrogen). Besides, the DNA quality was verified using a D-One spectrophotometer (Nanodrop). Multiplexed paired-end libraries (2 × 150 bp) were prepared using the Nextera XT DNA kit (Illumina, San Diego, CA, USA) and eventually sequenced with an Illumina NextSeq-500 instrument at a minimum of 50X coverage depth. Trimming and clipping were performed using AlienTrimmer v0.4.0 ^69^. Redundant or over-represented reads were reduced using the khmer software package v1.3 ^70^. Finally, sequencing errors were corrected using Musket v1.1 ^71^. A *de novo* assembly was performed for each strain using SPAdes v3.12.0 ^72^. The genomic sequences of the four reference strains were retrieved from public repositories (**Table S1**).

Additionally, the multidrug resistant isolate FRC0402 was subjected to long-read sequencing using Oxford Nanopore Technologies (ONT). Genomic DNA was extracted using the phenol-chloroform protocol combined with Phase Lock Gel tubes (Qiagen GmbH). Libraries were prepared using a 1D ligation sequencing kit (SQK-LSK-108) without fragmentation and sequenced using a MinION FLO-MIN-106 flow cell. Finally, ONT and Illumina short reads were combined to generate a hybrid assembly using Unicycler v0.4.4 (normal assembly mode, default parameters).

### Phylogeny, recombination and genomic sequence analyses

We built a core genome multiple sequence alignment (cg-MSA) from the assembled genome sequences. For this, the genome sequences were annotated using PROKKA v1.14.2 ^73^ with defaults parameters, resulting in GFF files. Roary v3.6 ^74^ was used to define protein-coding gene clusters, with a threshold set at 70% amino acid identity. Core genes were defined as being present in 95% of genomes and were concatenated into a cg-MSA by Roary. ClonalFrameML v1.11 ^75^ was used to build a phylogenetic tree based on the cg-MSA, which quantifies and accounts for the effects of recombination events. PhyML v20131022 ^76^ was used to build an initial tree.

We used Kleborate v1.0.0-beta (https://github.com/katholt/Kleborate), with the --resistance option, to identify (identity >80% and coverage >90%) known resistance genes in *C. diphtheriae* genomic sequences, based on the August 1, 2019 update of the ARG-Annot database. We used BLASTN (identity >80% and coverage >95%) to search for the presence of the *tox* gene and of genes associated with biovar Gravis (DIP351, DIP354 and DIP357) and for nitrate utilization (*narKGHJI)*.

MLST genotypes were defined using the international MLST scheme for *C. diphtheriae* and *C. ulcerans* ^12^.

### Genome wide association studies (GWAS)

The software treeWAS ^77^ was used to find genome-wide associations between either antimicrobial resistance phenotypes or biovar on the one hand, and genetic variants (both core-genome SNPs and accessory genome gene presence/absence) on the other hand. Core-genome SNPs were derived either from a mapping approach (Samtools v1.9 and GATK v3.4-0), which comprises intergenic regions; or from the alignment of core coding sequences found using Roary. We ran treeWAS v1.1 with default parameters, using as input the previously computed ClonalFrameML phylogenetic tree and distribution of homoplasies, in order to account for both the population structure and effect of recombination. For this analysis, susceptibility phenotypes were classified into resistant or susceptible categories based on zone diameter phenotypes using the CA-SFM/EUCAST 2019 cutoffs (**Figure 3, Table S5**). The seven chromosomal PBP coding sequences (including gene with locus tag RS14485 in RefSeq NC_002935.2, or DIP0637 in the original GenBank file) renamed by us as PBP4b) were extracted from the genomic sequences and translated into amino acid (AA) sequences, which were also analyzed for association with penicillin resistance.

### Cloning and transformation experiments

For ectopic expression in *C. glutamicum*, the *pbp2m* gene was amplified from *C. diphtheriae* strain FRC0402 and put under the control of the inducible *PgntK* promoter on the shuttle vector pTGR5 ^78^ **(Figure S10)** *. pbp2m* was assembled in this plasmid by Gibson assembly using the primers PBPdi_Fw (CAA AGA AAG GAT AAG ACC ATA TGA TGA CTA AGC ACA ATC GTT TCC GTC), PBPdi_Rv (TAC CTT AAG CGG CCG CTT TAT TGA ATT CCA GAG AAT TTC TGA ACA TCC G), pTGRdi_Fw (TAA AGC GGC CGC TTA AGG TAC C) and pTGRdi_Rv (ATG GTC TTA TCC TTT CTT TGG TGG CG).

*Escherichia coli* CopyCutter EPI400 (Lucigen) was used for cloning of the *pbp2m* gene and was grown in Luria-Bertani (LB) broth or agar plates at 37°C supplemented with 50 μg/ml kanamycin. The pTGR5_pbp2m plasmid was sequenced and electroporated into *Corynebacterium glutamicum* ATCC 13032. Positive colonies were grown in brain heart infusion (BHI) at 30°C and 120 rpm supplemented with 25 μg/ml kanamycin and 1% (w/v) gluconate when required for ectopic expression of Pbp2m.

### Mapping of SNPs on *C. diphtheriae* PBP sequences

Functional annotation of the different sequences was performed with InterPro ^79^. Conserved transpeptidation motifs SxxK, SxN and KTG were identified and mapped on the PBP sequences from *Corynebacteriales* based on the results of multiple sequence alignments performed with Clustal Omega ^80^. When uncertainty between a transmembrane domain and a signal peptide existed, a decision was made based on previous characterization of the homologous PBP in other *Corynebacteriales* in the literature.

## Data availability

The genomic sequencing data generated in this study were deposited in the European Nucleotide Archive (ENA) database and are accessible through the BioProject PRJEB22103.

## Code availability

No new code was used to analyse the findings in this study.

## Acknowledgements

We thank Vincent Enouf and the P2M core facility of Institut Pasteur for genomic sequencing. We are indebted to Collection de l’Institut Pasteur for providing reference strains of ribotypes, which were deposited following their described by Grimont and colleagues in 2004 ^40^. We thank Valerie Bouchez for help with Oxford Nanopore Technologies sequencing.

## Author contributions

S.B. conceived, designed and coordinated the study. A.C-L., E.B., M.B. and M.D. performed the microbiological cultures of the isolates and their biochemical and molecular characterizations. M.H., L.G.P., C.R., S.L.B., M.B.-P., M.D. and S.B. analysed the genomic and phenotypic data. J.T. reviewed the clinical source data of the isolates. Q.G. and A.-M.W. performed the pbp2m cloning experiments and analyzed the PBP sequences. X.D. provided help with the phylogenetic, recombination and GWAS analyses. S.B. wrote the initial version of the manuscript. All authors provided input to the manuscript and reviewed the final version.

## Supplementary Figures

**Figure S1. Biovar and *tox* status of the three strain subsets**

In the upper panel, the numbers of strains are given separately for the three subsets of strains; the recent clinical isolates one (right hand side) is broken down by individual year. Colors correspond to biovars (see key) and shaded areas denote *tox*-positive isolates. In the lower panel, the percentage of *tox*-positive strains (red bars) and of *tox*-positive or *tox*-negative strains per biovar (see key), are given for the entire dataset and for the three subsets separately; shaded sectors correspond to *tox*-positive strains within each biovar.

**Figure S2. Phylogenetic tree of *C. diphtheriae*, with isolates names**

The phylogenetic tree and outer information correspond to those in Figure 2, with the addition of isolates names, geographic origins and year of isolation.

**Figure S3. Genomic difference between reference strains PW8 (biovar Mitis) and NCTC13129 (Gravis)**

The genomic region of approx. 10 kb inserted in biovar Gravis strain NCTC13129 includes genes DIP0351, DIP0354 and *spuA* (DIP0357); these three accessory genes are strongly associated with biovar Gravis, as is the SNP at position 324,487.

**Figure S4. The distributions of zone diameter values for 19 antimicrobial agents, colored by the presence of the *tox* gene**

**A**: beta-lactams, macrolides and pristinamycin. **B**: other agents. X-axis: diameter in mm; Y-axis: number of strains. Colors inside the bars represent *tox*-positive isolates (red) or *tox*-negative isolates (grey). The three background colors represent the categorical interpretations according to EUCAST: resistant (salmon, left), intermediate (lighter salmon, middle) and susceptible (pale beige, right). The grey vertical bar corresponds to the proposed tentative ecological cutoff.

**Figure S5. The distributions of zone diameter values for 19 antimicrobial agents, colored by the main lineages (Mitis and Gravis)**

**A**: beta-lactams, macrolides and pristinamycin. **B**: other agents. X-axis: diameter in mm; Y-axis: number of strains. Colors inside the bars represent the two main lineages (see key in panel B). The three background colors represent the categorical interpretations according to EUCAST: resistant (salmon, left), intermediate (lighter salmon, middle) and susceptible (pale beige, right). The grey vertical bar corresponds to the proposed tentative ecological cutoff.

**Figure S6. Correlation plot of antimicrobial resistance phenotypes and genotypes.**

The correlation matrix between antimicrobial resistance genotype and phenotype is based on the correlation for binary variables (in the case of resistance genes: 1, presence; 0, absence; in the case of antimicrobial drugs: 1, resistant/intermediate; 0, susceptible) using the ‘corr.test’ function (Pearson method, which for a pair of binary variables equates to the Phi coefficient) from the ‘corrplot’ R package. Significant correlations were visualized utilizing the ‘corrplot’ function from the same package. Blank squares represent correlations without statistical significance (p > 0.05). Positive correlation is depicted by blue circles, whereas red circles represent significant negative correlation. The size and strength of color represent the numerical value of the Phi correlation coefficient. Black rectangles group genes commonly found together in the same strain. Genes *cmx* and *cmlA5* are known to be associated with chloramphenicol resistance, which was not tested here.

**Figure S7. treeWAS results plots for ciprofloxacin, rifampicin and trimethoprim**

Distribution of treeWAS scores obtained for genome-wide SNPs in association with ciprofloxacin, rifampicin and trimethoprim. Significant SNPs in *gyrA*, *rpoB* and *folA* are indicated.

**Figure S8. treeWAS results for amino acid polymorphisms in the chromosomal PBP coding genes of *C. diphtheriae***

Statistical significance of the treeWAS subsequent score obtained when testing the association of deduced amino-acid alterations in the seven chromosomal PBP sequences, and penicillin resistance phenotype. Within each of the seven panel the X-axis represent the amino acid sequence (numbers: AA positions), and the Y-axis the – log10(p-value). The positions of transglycosylase, transpeptidase, carboxypeptidase or other relevant domains of the PBP are shaded in grey. SNPs are represented as blue or orange circles (in alternance) at their corresponding position. The red bar indicates the 0.05 p-value position. The most significant SNP, at position 535 of PBP2b, is circled.

**Figure S9. Functional annotation and mapping of significant SNPs associated with *C. diphtheriae* chromosomal PBPs.**

Conserved transpeptidation motifs SxxK SxN KTG are indicated on the transpeptidase and carboxypeptidase domains by white lines. Significant SNPs associated with penicillin resistance are indicted by red pins. The PBPs correspond to the following genes: *pbp1a* (DIP2294), *pbp1b* (DIP0298), *pbp2a* (DIP0055), *pbp2b* (DIP1604), *pbp2c* (DIP1497), *pbp4* (DIP2005) and *pbp4b* (RS14485 = DIP0637). Pbp2m was not analyzed for amino-acid changes associated with penicillin resistance, as it corresponds to an accessory PBP.

**Figure S10. Construction strategy of plasmid pTGR5_pbp2m**

The *pbp2m* gene was PCR amplified and combined with plasmid pTGR5 using Gibson assembly as indicated.

**Figure S11. Genetic context of the pbp2m gene in *Corynebacterium***

The genomic context of *pbp2m* in *C. diphtheriae* and other *Corynebacterium* strains that possess this gene is given for representative genomes of the diversity that was found. Genes *pbp2m*, *blaB* and *lysR* are represented with a dark red background; these three genes were always associated and constitute the *pbp*-containing unit (PCU). Gene *ermX* is in yellow. A putative helicase often associated with the PCU is represented in pink; a relaxase gene is shaded in green. Black arrows represent insertion sequence genes. Six groups were defined based on conserved features, as indicated. Dark grey parallelepiped joining different genomes represent homology levels, as indicated in the gradient key. Strains of the present study with identical structures as those represented are indicated in parentheses below the strain name of the representative genome. The scale bar represents 12 kb.

