## Supplementary figures and images for "Population genomics and antimicrobial resistance in *Corynebacterium diphtheriae*"

### FigureS1

#### Slide 1
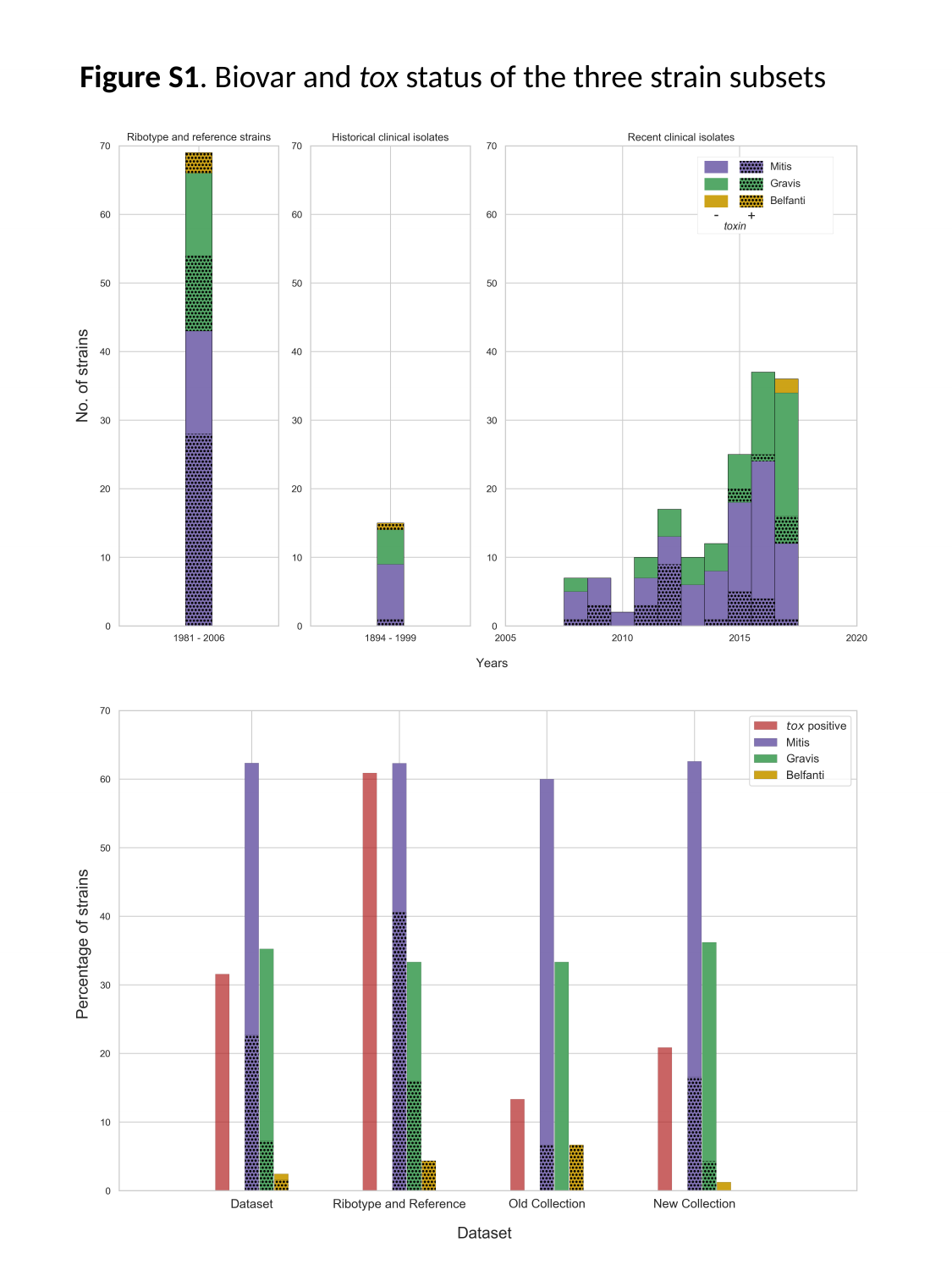

Figure S1. Biovar and tox status of the three strain subsets

### FigureS2

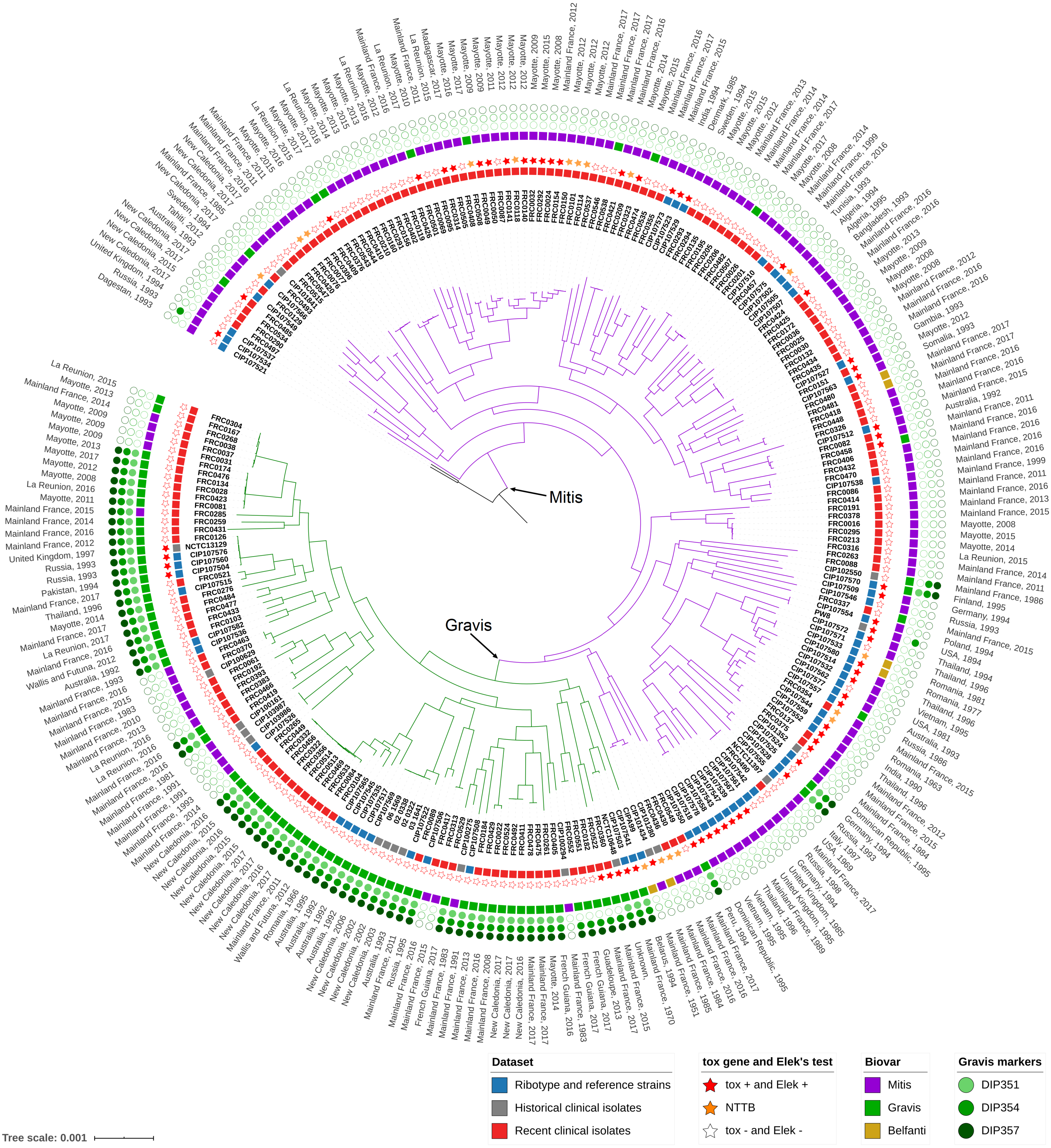

### FigureS4a

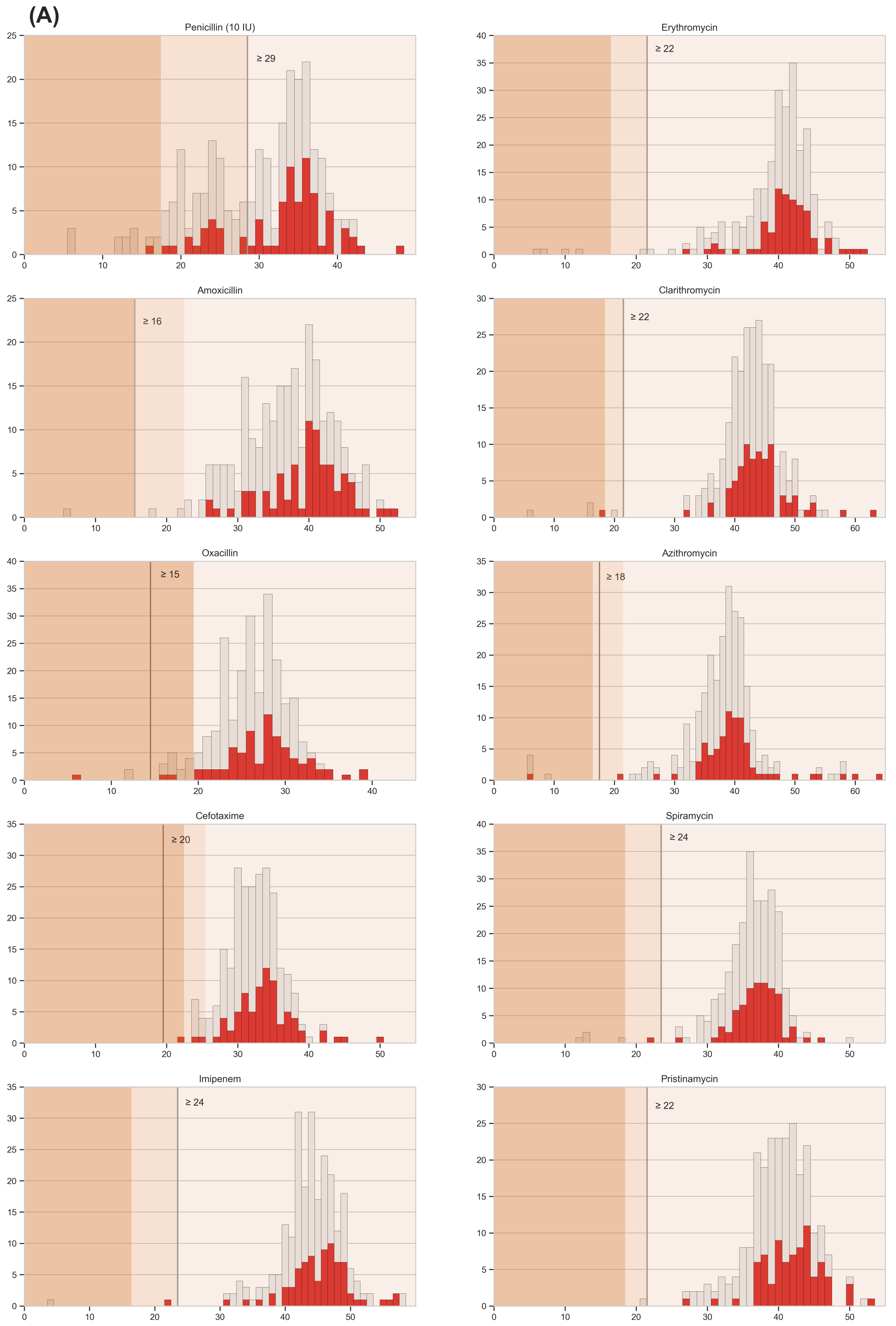

### FigureS4b

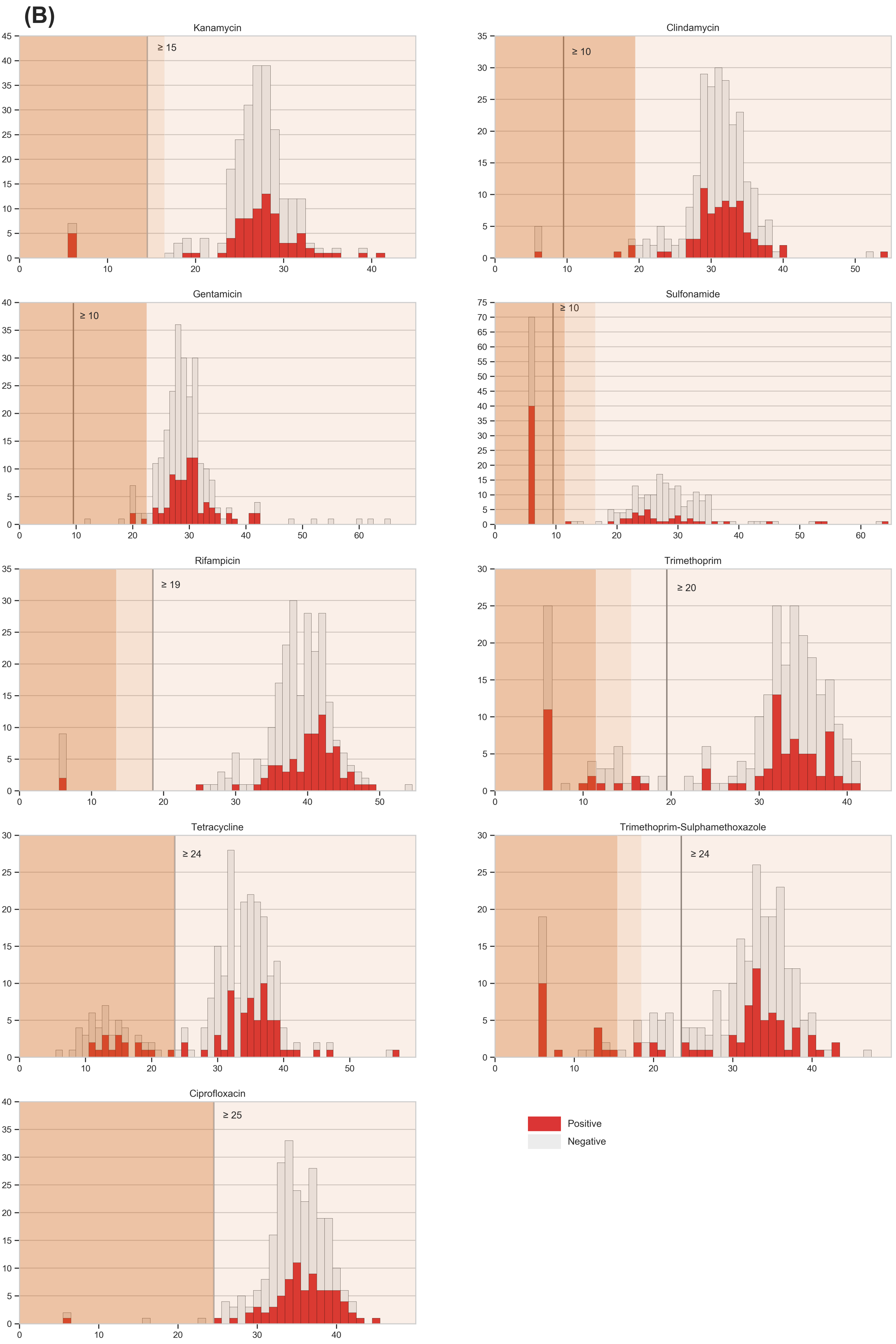

### FigureS5a

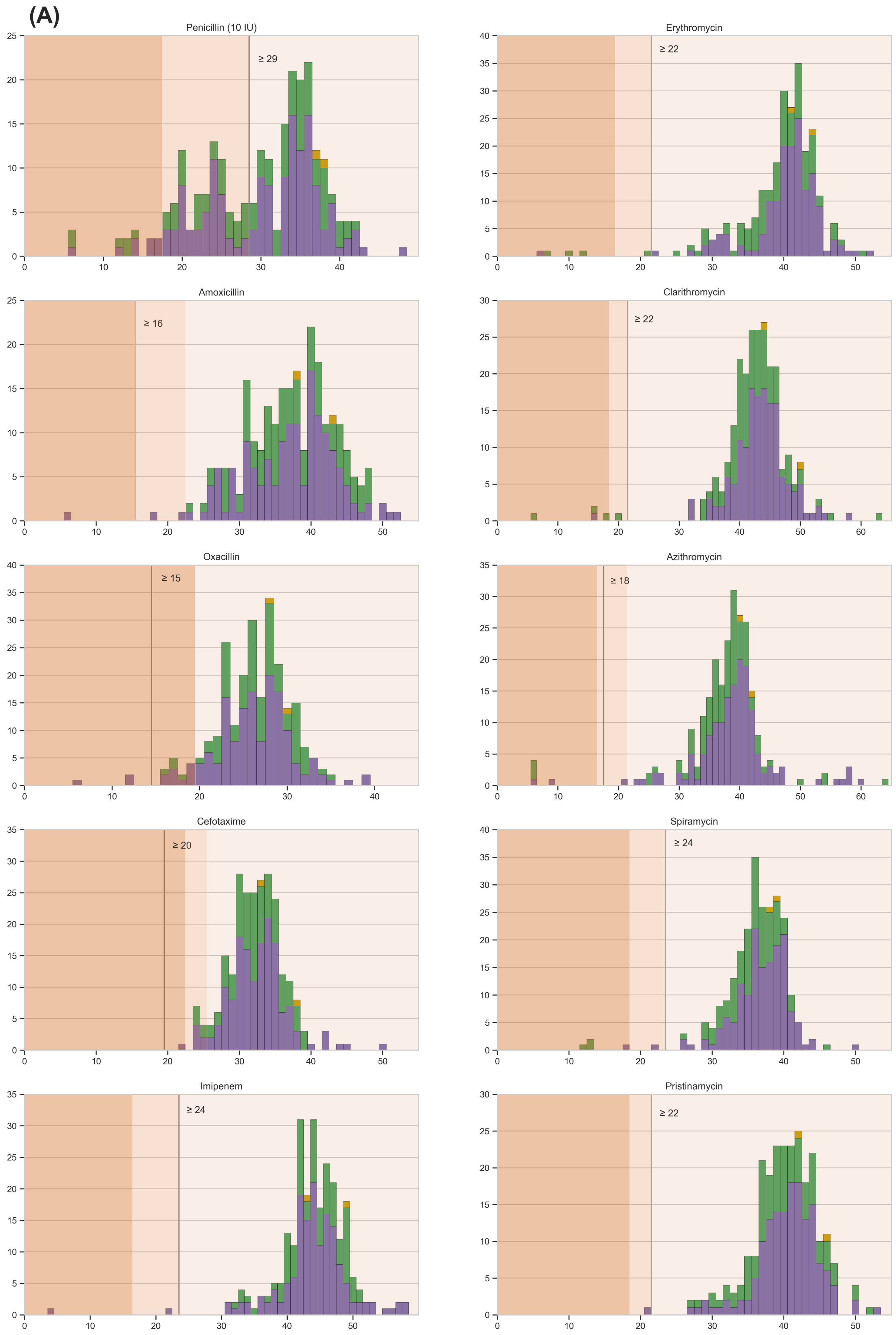

### FigureS5b

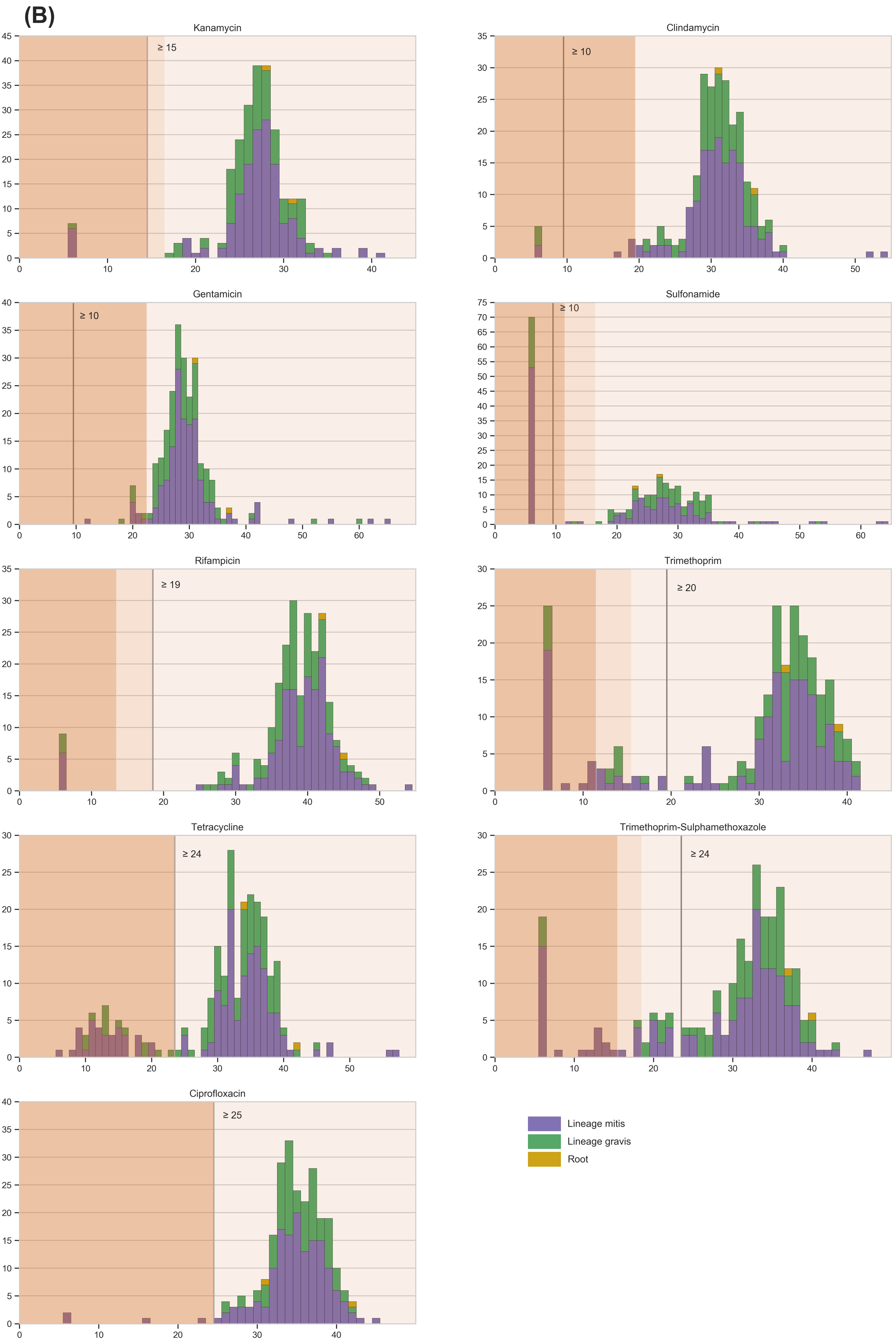

### FigureS8

#### Slide 1
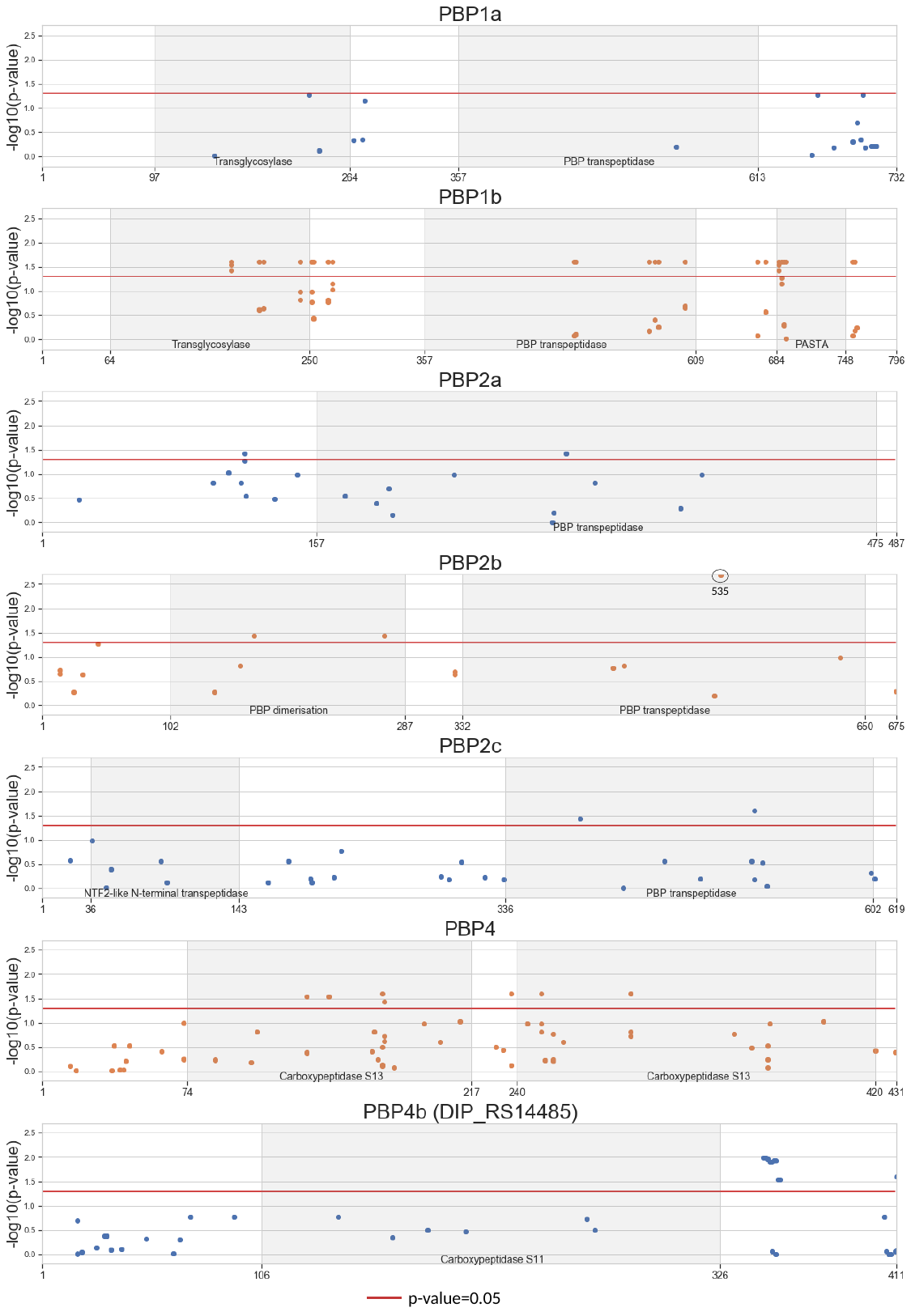

535
p-value=0.05

### FigureS10

#### Slide 1
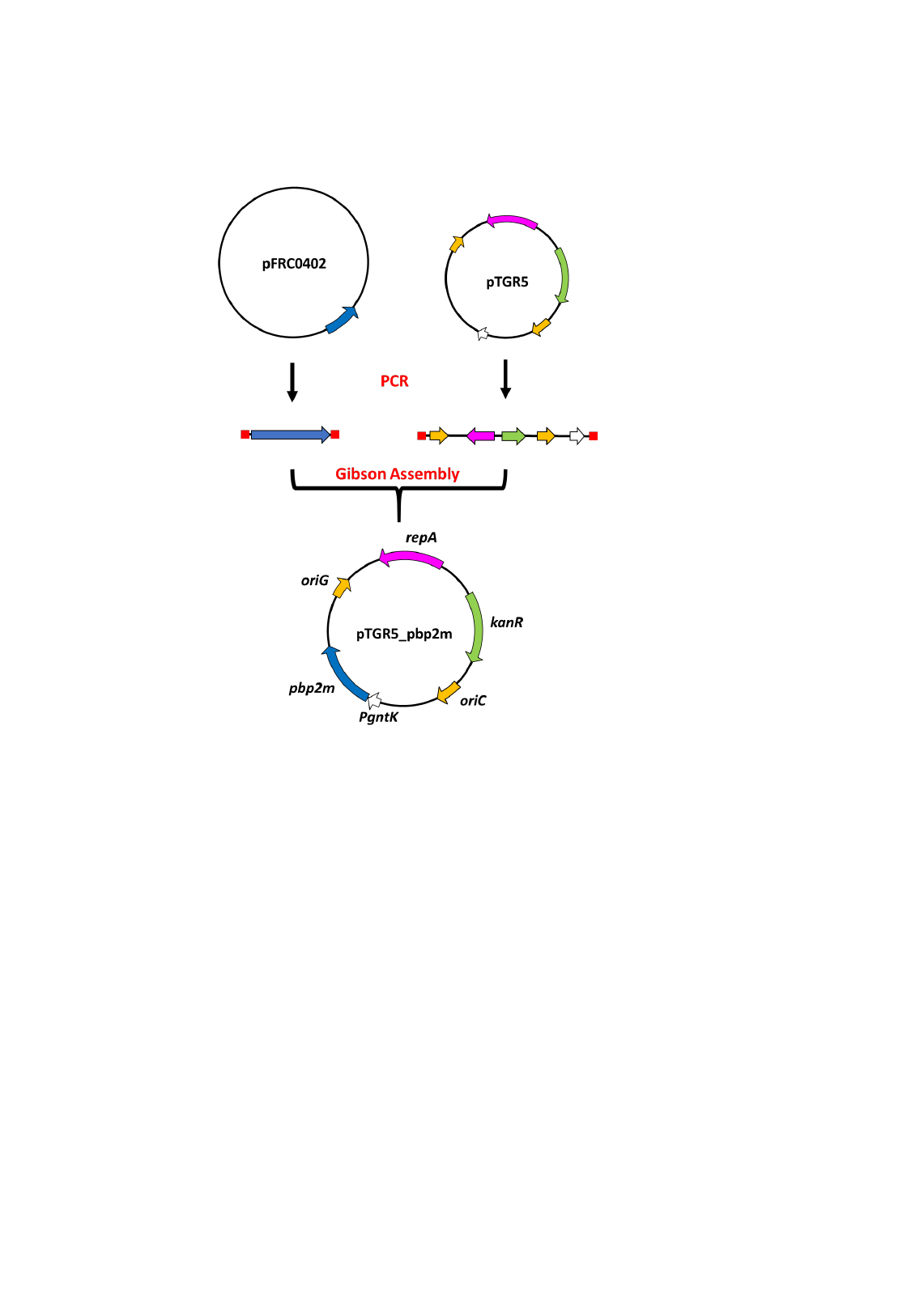
