## Supplementary material for "Population genomics and antimicrobial resistance in *Corynebacterium diphtheriae*": FigureS3

### Slide 1
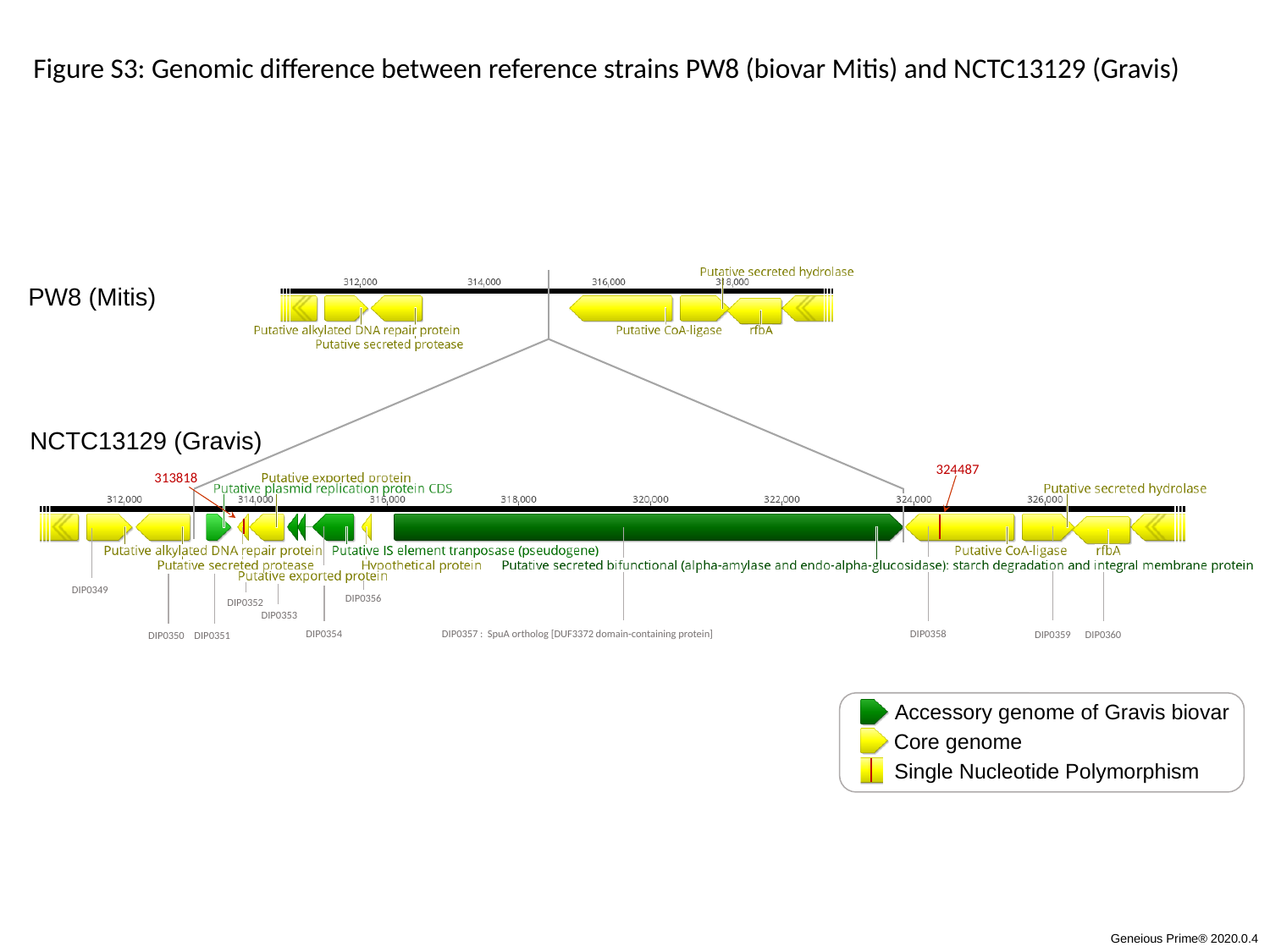

Figure S3: Genomic difference between reference strains PW8 (biovar Mitis) and NCTC13129 (Gravis)
PW8 (Mitis)
NCTC13129 (Gravis)
324487
313818
DIP0349
DIP0356
DIP0352
DIP0353
DIP0354
DIP0357 : SpuA ortholog [DUF3372 domain-containing protein]
DIP0358
DIP0359
DIP0360
DIP0351
DIP0350
Accessory genome of Gravis biovar
Core genome
Single Nucleotide Polymorphism
Geneious Prime® 2020.0.4
