## Supplementary material for "Population genomics and antimicrobial resistance in *Corynebacterium diphtheriae*": FigureS6

### Slide 1
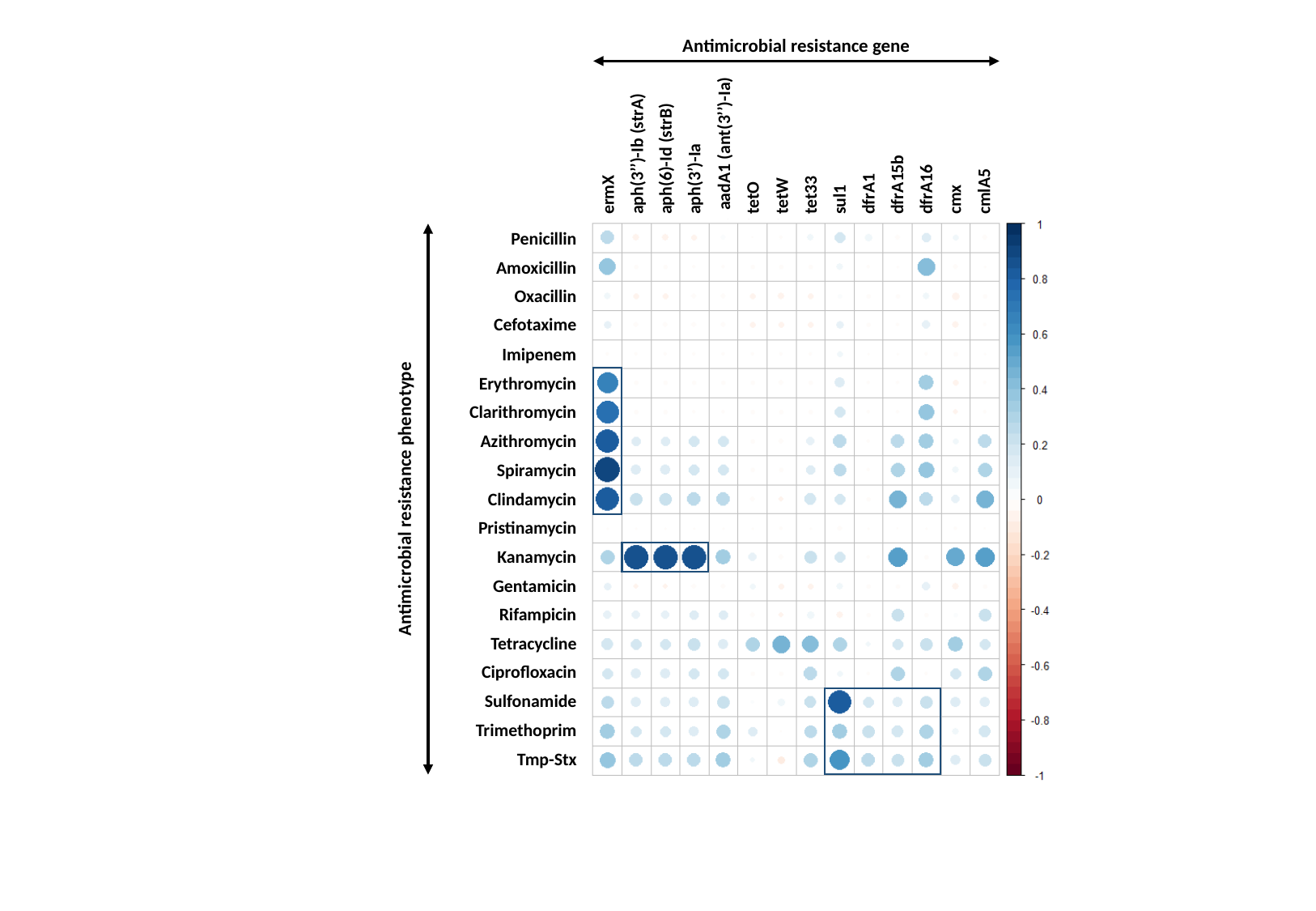

Antimicrobial resistance gene
ermX
aph(6)-Id (strB)
 aadA1 (ant(3’’)-Ia)
tetW
sul1
dfrA15b
cmx
aph(3’’)-Ib (strA)
aph(3’)-Ia
tetO
tet33
dfrA1
dfrA16
cmlA5
Penicillin
Oxacillin
Imipenem
Clarithromycin
Spiramycin
Pristinamycin
Gentamicin
Amoxicillin
Cefotaxime
Erythromycin
Azithromycin
Clindamycin
Kanamycin
Rifampicin
Tetracycline
Ciprofloxacin
Sulfonamide
Trimethoprim
Tmp-Stx
Antimicrobial resistance phenotype
