## Supplementary material for "Population genomics and antimicrobial resistance in *Corynebacterium diphtheriae*": FigureS7

### Fluoroquinolone

#### Ciprofloxacin

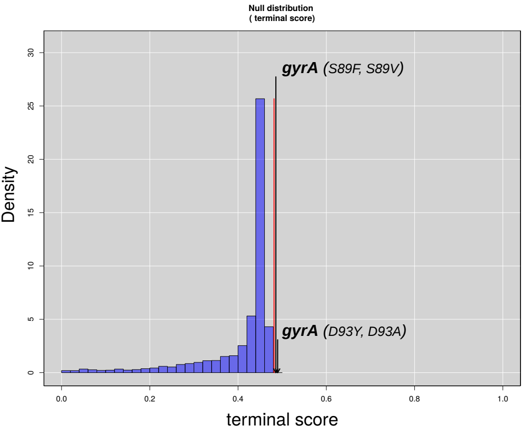


### RNA metabolism

#### Rifampicin

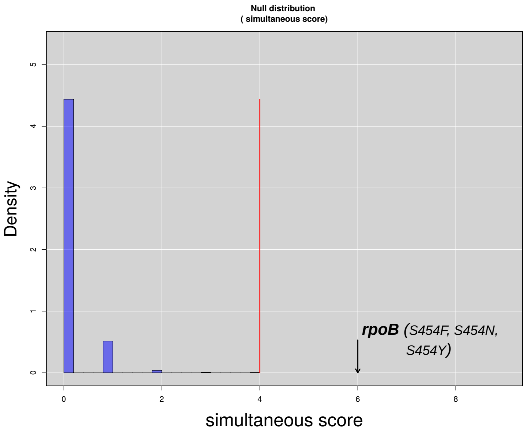


### Folic acid metabolism

#### Trimethropin

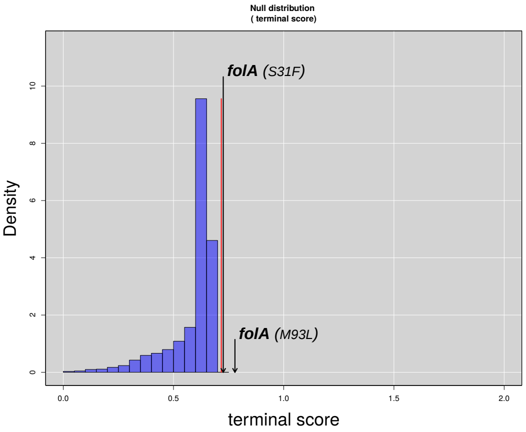
