## Supplementary material for "Population genomics and antimicrobial resistance in *Corynebacterium diphtheriae*": FigureS9

### Slide 1
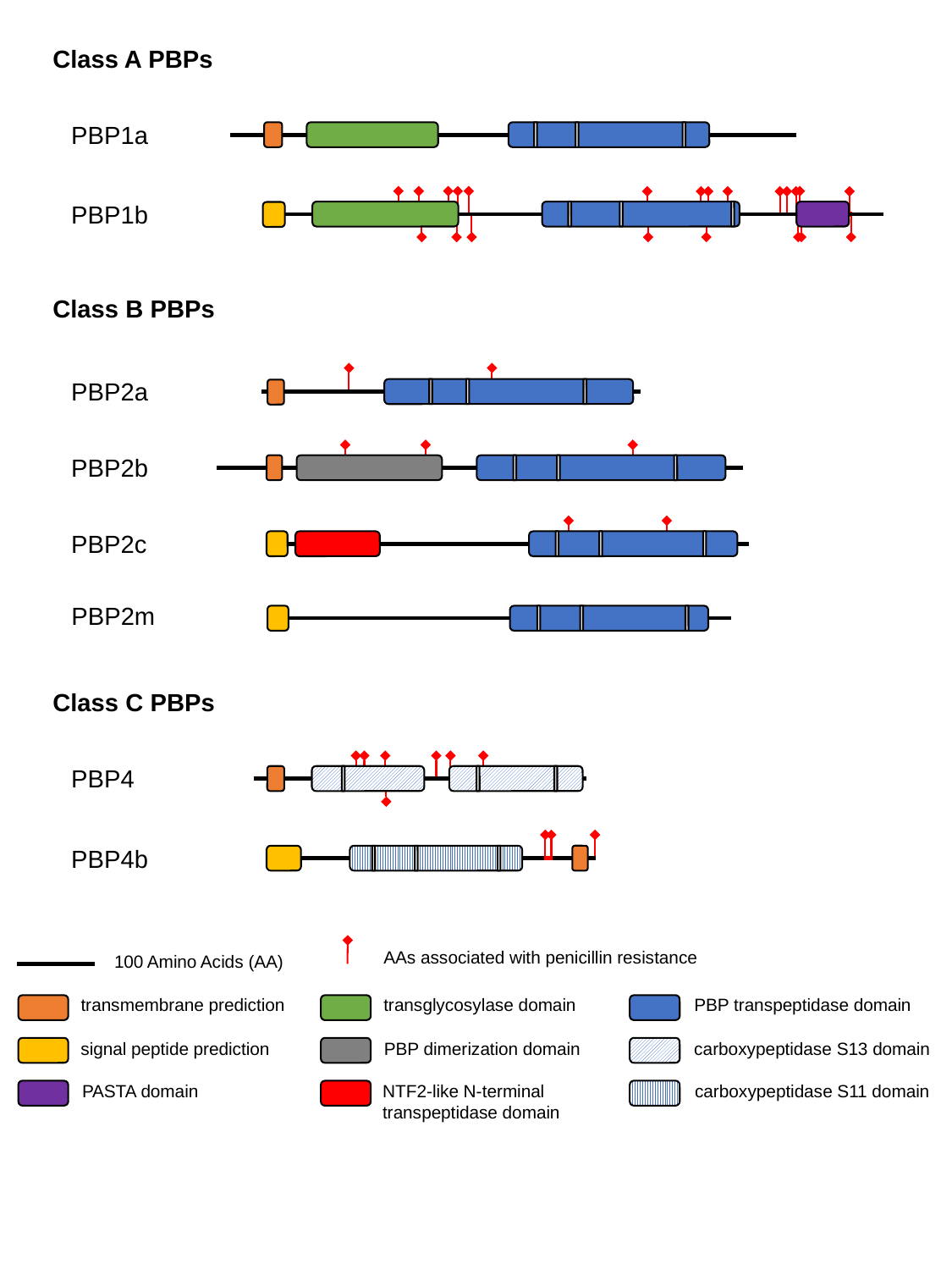

Class A PBPs
PBP1a
PBP1b
Class B PBPs
PBP2a
PBP2b
PBP2c
PBP2m
Class C PBPs
PBP4
PBP4b
AAs associated with penicillin resistance
100 Amino Acids (AA)
transmembrane prediction
transglycosylase domain
PBP transpeptidase domain
signal peptide prediction
PBP dimerization domain
carboxypeptidase S13 domain
PASTA domain
NTF2-like N-terminal transpeptidase domain
carboxypeptidase S11 domain
