## Supplementary material for "Population genomics and antimicrobial resistance in *Corynebacterium diphtheriae*": FigureS11

### Slide 1
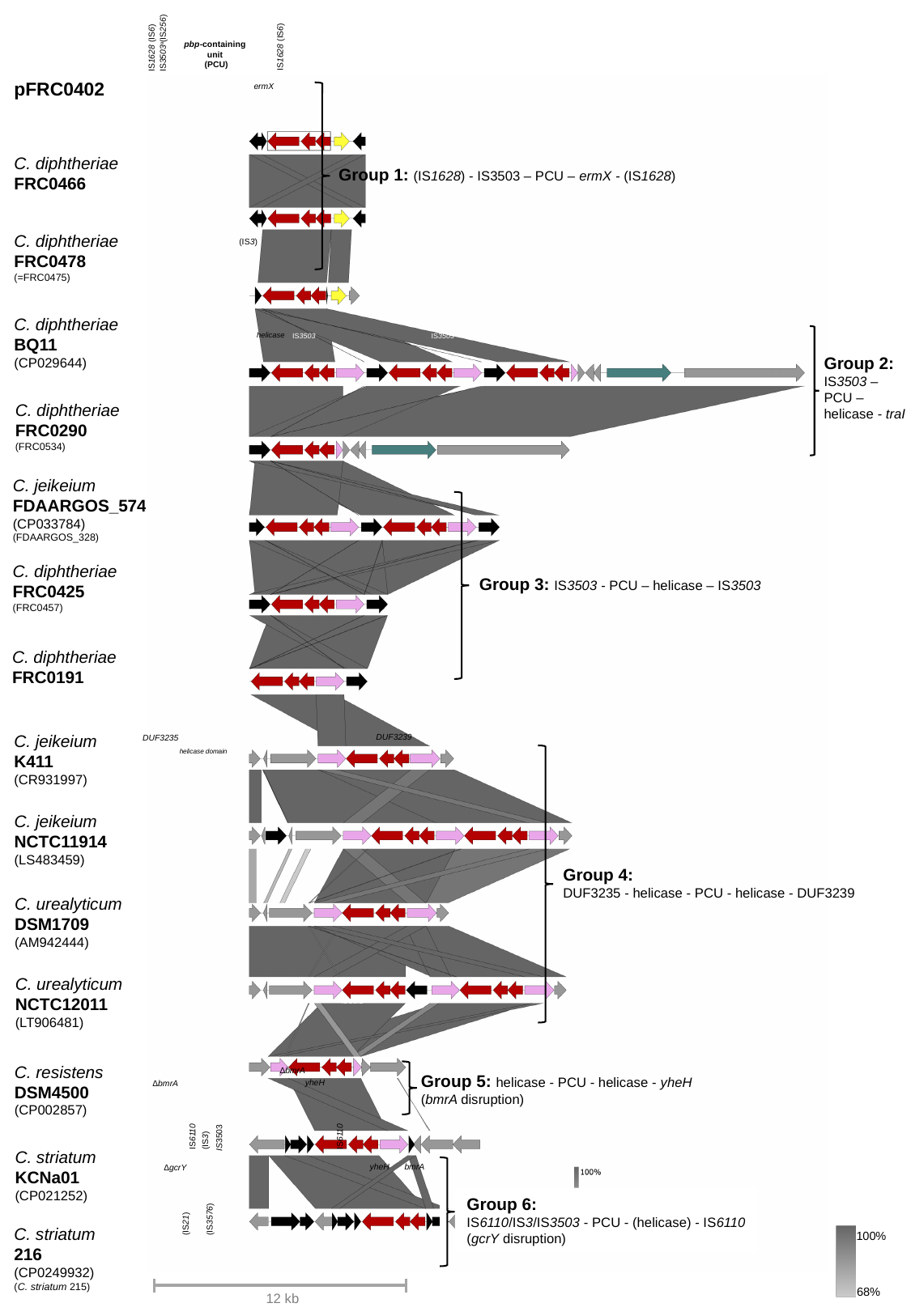

IS3503^a^(IS256)
IS1628 (IS6)
IS1628 (IS6)
pbp-containing
unit
(PCU)
pbp2m
ermX
lysR
blaB
(IS3)
traI (relaxase)
helicase
IS3503
IS3503
IS3503
DUF3239
DUF3235
helicase domain
IS3503
ISCur2
∆bmrA
yheH
∆bmrA
IS6110
IS6110
IS3503
(IS3)
yheH
bmrA
∆gcrY
(IS3576)
(IS21)
Group 1: (IS1628) - IS3503 – PCU – ermX - (IS1628)
Group 2:
IS3503 –
PCU –
helicase - traI
Group 3: IS3503 - PCU – helicase – IS3503
Group 4:
DUF3235 - helicase - PCU - helicase - DUF3239
Group 5: helicase - PCU - helicase - yheH
(bmrA disruption)
Group 6:
IS6110/IS3/IS3503 - PCU - (helicase) - IS6110
(gcrY disruption)
100%
68%
12 kb
pFRC0402
C. diphtheriae
FRC0466
C. diphtheriae
FRC0478
(=FRC0475)
C. diphtheriae
BQ11
(CP029644)
C. diphtheriae
FRC0290
(FRC0534)
C. jeikeium
FDAARGOS_574
(CP033784)
(FDAARGOS_328)
C. diphtheriae
FRC0425
(FRC0457)
C. diphtheriae
FRC0191
C. jeikeium
K411
(CR931997)
C. jeikeium
NCTC11914
(LS483459)
C. urealyticum
DSM1709
(AM942444)
C. urealyticum
NCTC12011
(LT906481)
C. resistens
DSM4500
(CP002857)
C. striatum
KCNa01
(CP021252)
C. striatum
216
(CP0249932)
(C. striatum 215)
